# Integrated metabolomic and metagenomic profiling reveals distinct signatures in treatment-naïve multiple sclerosis patients

**DOI:** 10.64898/2025.11.30.691404

**Authors:** Jessica Perego, Valeria Mannella, Laura Manuto, Mireia Valles-Colomer, Francesco Asnicar, Elisa Piperni, Albert Garcia-Valiente, Marcella Bonanomi, Daniela Gaglio, Federico Montini, Vittorio Martinelli, Nicola Segata, Gianvito Martino, Paola Panina-Bordignon

## Abstract

Imbalances in gut microbiota composition and microbiota-associated metabolites have been linked with several neurological diseases, including multiple sclerosis (MS). However, a comprehensive multi-biofluid approach remains lacking. Most studies on MS include patients already receiving treatment and often neglect to account for sex-related differences, which could serve as potential confounding factors. Moreover, they mostly investigate only stools or plasma metabolomics. In this study, we recruited 18 treatment-naïve neuroinflammatory patients at diagnosis and compared them with 20 healthy controls, matched for sex and age. We conducted multi-biofluids metabolomic analysis of urine, stool, serum, and cerebrospinal fluid, complemented by taxonomical and functional profiling of the gut microbiome using shotgun metagenomic sequencing. Our results show that MS patients exhibit distinct microbiome composition and urinary metabolomic profiles compared to healthy controls. Furthermore, neuroinflammation is associated with dysregulation of microbially-produced short-chain fatty acids, their intestinal absorption and systemic bioavailability, as illustrated by the altered plasma levels.

## INTRODUCTION

Multiple sclerosis (MS) is an inflammatory, demyelinating disease of the central nervous system (CNS) in which both genetic and environmental factors contribute to disease onset¹. However, the low concordance rate in monozygotic twins highlights the substantial role of environmental triggers in MS susceptibility^1^. Recently, the gut microbiota and its microbial metabolites have emerged as key environmental determinants in the pathobiology of a wide range of neurological conditions³, including MS⁴. Not only the gut microbiome but also its metabolic pathways and circulating gut-derived metabolites are dysregulated in people with MS (PwMS)^2,3^. Although it remains unclear whether these microbial alterations are a cause or a consequence of the disease, we sought to examine patients as close as possible to disease onset by analyzing individuals at diagnosis, before treatment initiation. This approach provided two major advantages: (1) it eliminated treatment as a significant confounding factor, as all patients were treatment-naïve; and (2) it enabled us to analyze cerebrospinal fluid (CSF), which was collected for diagnostic purposes.

Studies in mice and humans suggest that the microbiota can influence the onset and progression of disease through immune effector cells as well as soluble metabolic, immune, and neuroendocrine factors modulated by gut microbial activity^4,5^. While microbial changes in MS have been reported, most published studies focus on patients already undergoing treatment, which can significantly influence both gut microbiota composition and metabolomic profiles, potentially confounding the interpretation of disease-specific changes^3^. Indeed, many commonly used drugs possess antimicrobial properties or induce substantial shifts in the gut microbiome, raising the possibility that therapeutic efficacy may, in part, be mediated by microbiome modulation^6–8^. Notably, pharmacological interventions can alter the gut microbiota, resulting in both therapeutic benefits and potential adverse effects. Furthermore, most published studies focus mainly on taxonomic characterization of the bacterial communities, often neglecting other key components of the gut microbiome, such as viruses, fungi, and other eukaryotic organisms. In addition, functional metabolomic profiling is typically limited to stool or plasma samples, offering only a limited perspective on the systemic metabolomic alterations associated with disease. Previous studies suggest that PwMS exhibit metabolic alterations in their systemic metabolomic profiles^3^, consistent with findings from metabolomics studies in other neurological disorders, such as Alzheimer’s^9,10^ and Parkinson’s diseases^11,12^. While extensive research has investigated gut microbiota dysbiosis in MS and its association with immune modulation and disease progression, no studies to date have specifically integrated gut microbiota composition with urinary, stool, and plasma metabolomic profiles from the same cohort. As a result, comprehensive data on the microbial influence across multiple biofluids in newly diagnosed, treatment-naïve PwMS, particularly regarding sex-dependent features, are still lacking. Our study addresses this gap by performing a sex-specific comparison between newly diagnosed, treatment-naïve PwMS and healthy controls (HC), examining both stool microbiota composition and multi-fluid metabolomic profiles across urine, stool, plasma, and CSF

Among the key microbiota-synthesized metabolites that have been linked to neurological disorders^13^ in general, and MS^14^ in particular, short-chain fatty acids (SCFAs) have been strongly associated with disease onset and progression. SCFAs are a group of fatty acids with two to six carbon atoms, produced by the gut microbiota primarily through the fermentation of indigestible dietary fibers and resistant starch. The three most physiologically relevant SCFAs are acetate, propionate, and butyrate. Short-chain fatty acids (SCFAs) are key regulators of gut health, immune modulation, and various metabolic processes, and have been associated with anti-inflammatory effects. Their potential role in MS has generated increasing attention due to these anti-inflammatory properties. However, although several studies have explored the relationship between SCFAs and MS, none have assessed SCFA levels across multiple biofluids in a treatment-naïve cohort.

## RESULTS

### Cohort of treatment-naïve, newly diagnosed people with MS

A total of 18 patients with neuroinflammatory conditions and 20 genetically unrelated HC were recruited between November 2021 and November 2022 at San Raffaele Hospital, Milan (Italy) (Table 1). Among the 18 patients, 17 had a confirmed diagnosis of MS, while one had clinical symptoms suggestive of MS (*i.e*., clinically isolated syndrome) but failed to confirm the MS diagnosis. Among the 17 PwMS, a single patient was diagnosed with primary progressive MS (PMS), while 16 were diagnosed with relapsing-remitting MS (RRMS). Considering that, statistically, RRMS affects 85–90% of patients at diagnosis, while PMS occurs in 10–15% of PwMS^15^, our cohort reflects the predominance of the RRMS form at diagnosis. The mean disease duration was 5.2 years for women and 1.6 years for men (Table 1), a difference driven mainly by a female patient with a 21-year disease duration (Fig. S1), though this was not statistically significant (Mann-Whitney test, p = 0.4477). The disease duration has been calculated by the neurologists starting from the earliest possible symptoms of MS. The long disease duration might be because symptoms were mild and neglected by the patient or confused with an improper diagnosis. The mean Expanded Disability Status Scale (EDSS)^16^ score for PwMS was 1.7 with no substantial difference between sexes (Table 1). Since the only patient without a confirmed MS diagnosis had clinical and experimental features consistent with the cohort, we included this individual in the MS group for all analyses. All the samples were collected at the diagnosis, before the initiation of disease-modifying therapies (DMTs). Our cohort exhibited a nearly 1:1 female - to - male ratio in MS and HC, allowing us to perform sex-specific analyses. Participants had an average of 37 years, with HC being slightly but not significantly younger (Table 1 and Fig. S1). Most donors had a normal body mass index (BMI), though male PwMS showed a marginally higher BMI compared to male HC and female PwMS (one-way ANOVA, p-value < 0.05, Table 1, Fig. S1). All patients completed a clinical survey reporting disease status and clinical parameters (Table S1), and all participants completed a subject survey detailing medication use, lifestyle, and pre-existing extra-intestinal and non-neurological conditions (Table 1). Participants were not on a standardized diet but followed their usual dietary regimen.

**Table 1.**
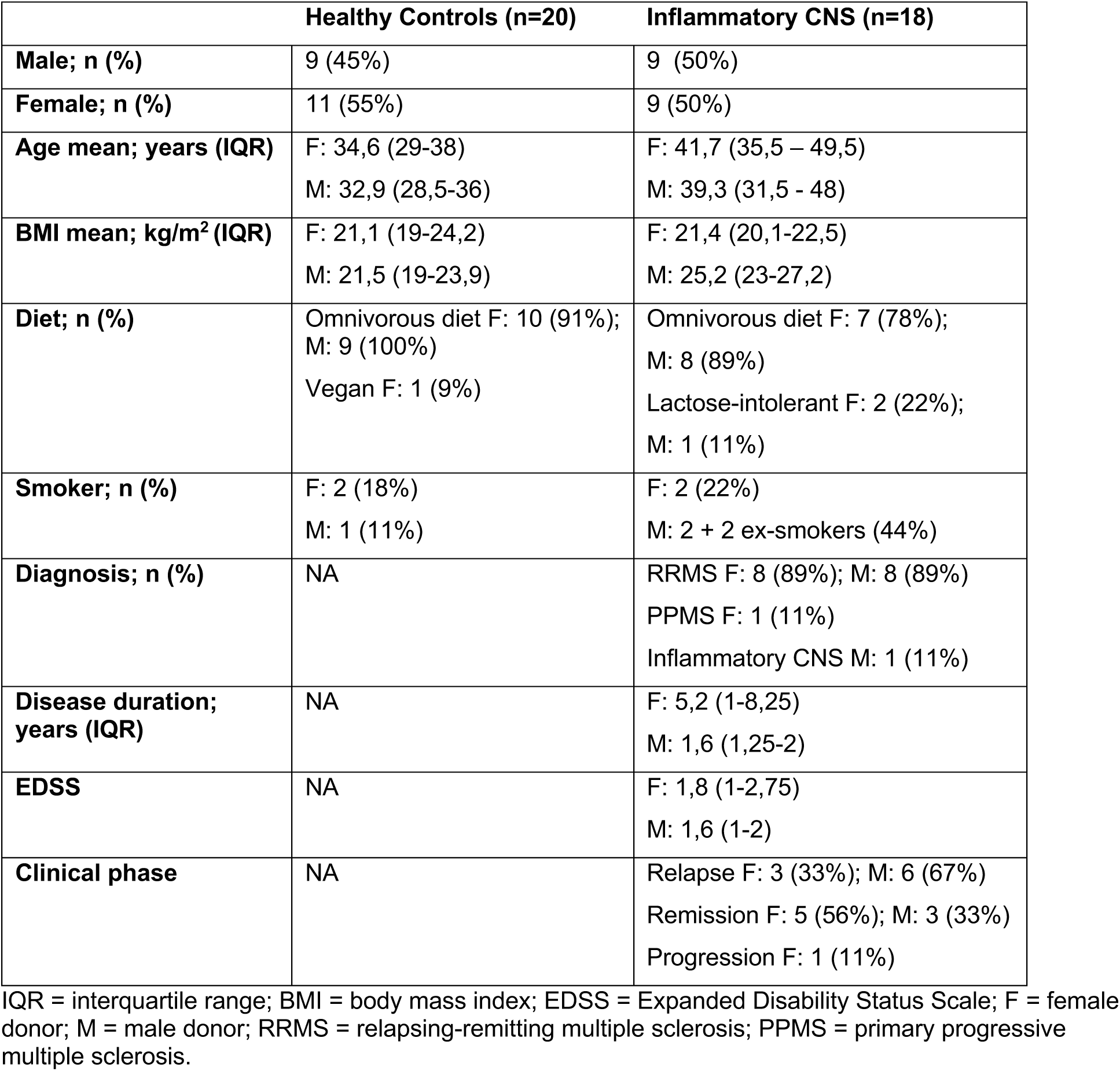
Cohort characterization.

|  | Healthy Controls (n=20) | Inflammatory CNS (n=18) |
| --- | --- | --- |
| **Male; n (%)** | 9 (45%) | 9 (50%) |
| **Female; n (%)** | 11 (55%) | 9 (50%) |
| **Age mean; years (IQR)** | F: 34,6 (29-38)<br>M: 32,9 (28,5-36) | F: 41,7 (35,5 – 49,5)<br>M: 39,3 (31,5 - 48) |
| **BMI mean; kg/m<sup>2</sup> (IQR)** | F: 21,1 (19-24,2)<br>M: 21,5 (19-23,9) | F: 21,4 (20,1-22,5)<br>M: 25,2 (23-27,2) |
| **Diet; n (%)** | Omnivorous diet F: 10 (91%);<br>M: 9 (100%)<br>Vegan F: 1 (9%) | Omnivorous diet F: 7 (78%);<br>M: 8 (89%)<br>Lactose-intolerant F: 2 (22%);<br>M: 1 (11%) |
| **Smoker; n (%)** | F: 2 (18%)<br>M: 1 (11%) | F: 2 (22%)<br>M: 2 + 2 ex-smokers (44%) |
| **Diagnosis; n (%)** | NA | RRMS F: 8 (89%); M: 8 (89%)<br>PPMS F: 1 (11%)<br>Inflammatory CNS M: 1 (11%) |
| **Disease duration; years (IQR)** | NA | F: 5,2 (1-8,25)<br>M: 1,6 (1,25-2) |
| **EDSS** | NA | F: 1,8 (1-2,75)<br>M: 1,6 (1-2) |
| **Clinical phase** | NA | Relapse F: 3 (33%); M: 6 (67%)<br>Remission F: 5 (56%); M: 3 (33%)<br>Progression F: 1 (11%) |

IQR = interquartile range; BMI = body mass index; EDSS = Expanded Disability Status Scale; F = female donor; M = male donor; RRMS = relapsing-remitting multiple sclerosis; PPMS = primary progressive multiple sclerosis.

### Shotgun metagenomic analysis suggests a decreased alpha diversity in female PwMS

Alpha diversity was measured using several indices: Observed, Simpson, Fisher, and Pielou (Fig. 1A-B). No significant differences in alpha diversity were observed between MS and HC groups (Wilcoxon Rank-Sum test, p < 0.05, FDR < 0.1; Fig. 1B), as also reported in other independent cohorts^1,3,17,18^. However, PwMS, particularly female donors, tended to have lower alpha diversity compared to female HC.

**Figure 1.**
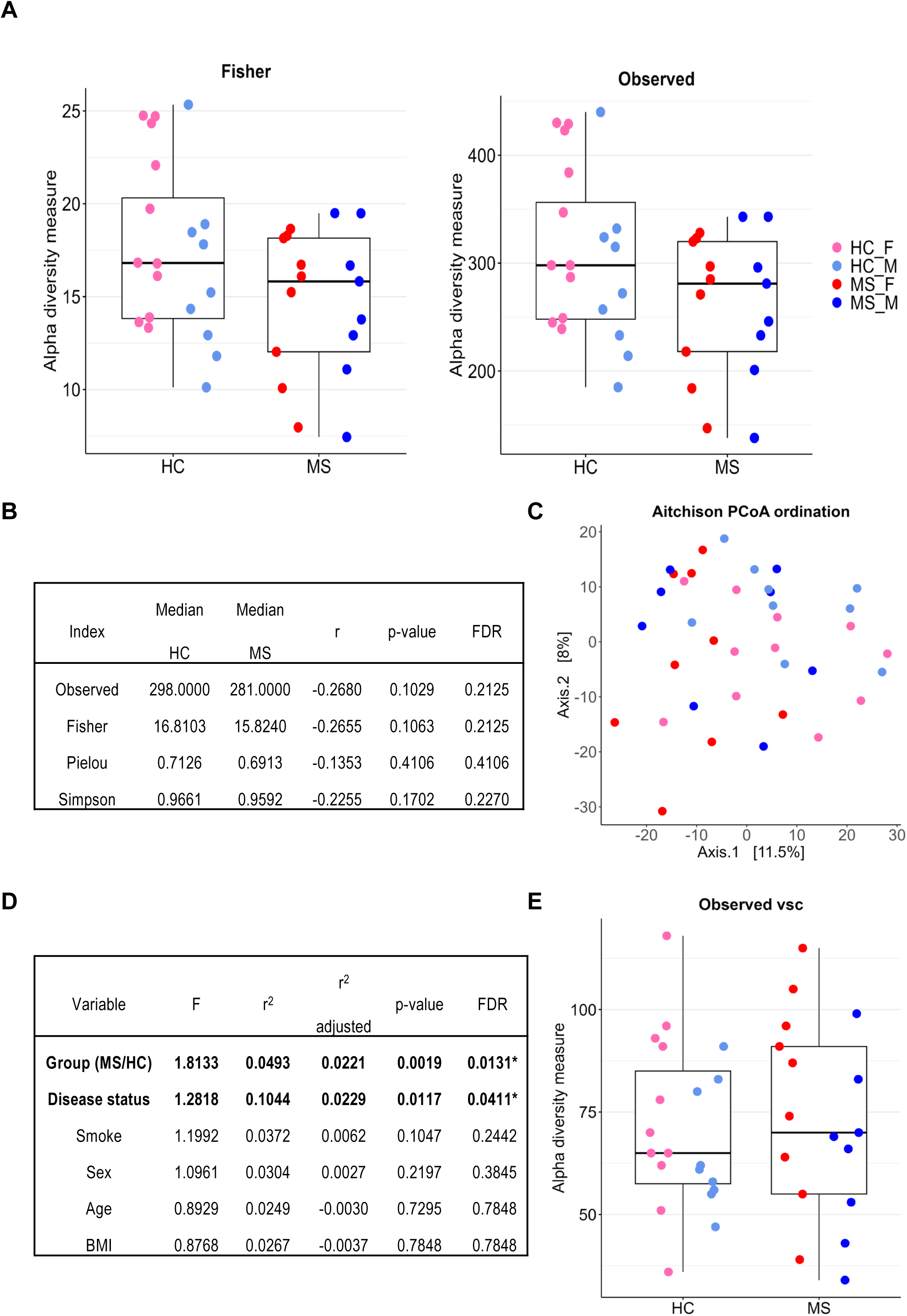
Shotgun metagenomic analysis suggests an altered bacterial and viral alpha diversity in female PwMS. (A) Bar plot of the bacterial alpha diversity indexes, Fisher and Observed, median + interquartile range (IQR). Colored dots represent the sex component (pink = female HC; light blue = male HC; red = female MS; dark blue = male MS). Wilcoxon rank-sum test, p-value <0,05, FDR <0,1. (B) Statistics on bacterial alpha diversity are calculated with multiple indexes: Observed, Simpson, Fisher, and Pielou. r = effect size. Wilcoxon rank-sum test, p-value <0,05, FDR <0,1. (C) PCoA of Aitchison distance between all samples considering the bacterial microbiota composition. Colored dots represent both the sex component and the group category (MS or HC). PERMANOVA FDR < 0.05. (D) Statistics on the samples’ variables correlation with the bacterial microbiota variability. F = F score, r^2^ = coefficient of determination; PERMANOVA FDR <0,1. Variables in bold are statistically significant (* FDR <0,1). (E) Bar plot of the observed Viral Sequence Clusters (VSCs), median + IQR. Colored dots represent the sex component. Median HC=65; median MS=70. p-value = 0.68, calculated with the Wilcoxon rank sum test. (A-E) HC n=20; MS n= 17.

Beta diversity analysis showed no clustering separation between MS and HC, as illustrated by the principal coordinates analysis (PCoA) based on Aitchison distance (Fig. 1C). Nonetheless, group identity (HC or MS) and disease status (“relapse,” “remission,” “progression,” or “healthy control”) significantly correlated with microbiota interindividual variation (PERMANOVA FDR < 0.1; Fig. 1D), while age, sex, BMI, and smoking did not.

Although most of the studies consider only the bacterial component, the complexity of the microbiome is given by the co-existence of bacterial, viral, fungal, and protozoan species. We thus dug deeper into the metagenomic data to capture this additional layer of microbial diversity (Fig. 1E). Although we could not find a statistically significant difference due to the limited cohort size, we still noted a trend of increased virome alpha diversity in PwMS compared to HC (Fig. 1E). At the species level, we found that the viral species M1098 is detected in all MS patients but only in 60% of HC (Fig. S2A-B). On the other hand, the species M311 is absent in 94% of PwMS, but present in 35% of HC (Fig. S2A-B). Unfortunately, these species still lack a detailed description, so we cannot speculate further. As a general remark, most of the viruses differently expressed between PwMS and HC and reported in Fig. S2B, except M311, seem to be present with a higher frequency in PwMS.

The last microbiome component we considered was the presence of Blastocystis in our samples. We selected *Blastocystis* to investigate specifically the microeukaryotic component of the microbiota since it has been recently reported by Piperni et al^19^ that the presence of this unicellular eukaryote defined a specific bacterial signature and was positively associated with more favorable cardiometabolic profiles and negatively with obesity and disorders linked to altered gut ecology. Different *Blastocystis* subtypes (ST 1, 3, 6, and 9) were identified in only four samples, all belonging to the HC group. The positive samples were equally distributed to male and female donors, and the positive donors had an age gap of 16 years (33-49), suggesting that the features are neither sex nor age dependent (Fig. S2B). To our knowledge, no previous study assessed the presence of *Blastocystis* in the metagenome of PwMS.

### Microbial taxonomic profile of PwMS reveals a decrease in butyrate-producing bacteria

We subsequently leveraged metagenomic data to characterize changes in the core-microbiota taxa and pathways profiles in PwMS compared to HC. Concerning the core microbiota taxa, *i.e.,* those with a minimum abundance of 0.001 and a prevalence in more than 80% of subjects, our study confirmed MS-driven microbiota alterations in line with previously published studies. At the *phylum* level, PwMS showed a statistically significant decrease in Firmicutes^18,20–22^ and an increase in Actinobacteria^23^ (Fig. 2A), with a p-value of 0.0658 after FDR correction. Firmicutes, primarily Gram-positive, are essential for fermenting complex carbohydrates into short-chain fatty acids (SCFAs), notably butyrate^24–26^. Actinobacteria, increased in PwMS, are known to influence the gut-brain axis via neuroactive metabolites such as gamma-aminobutyric acid (GABA)^27^.

**Figure 2.**
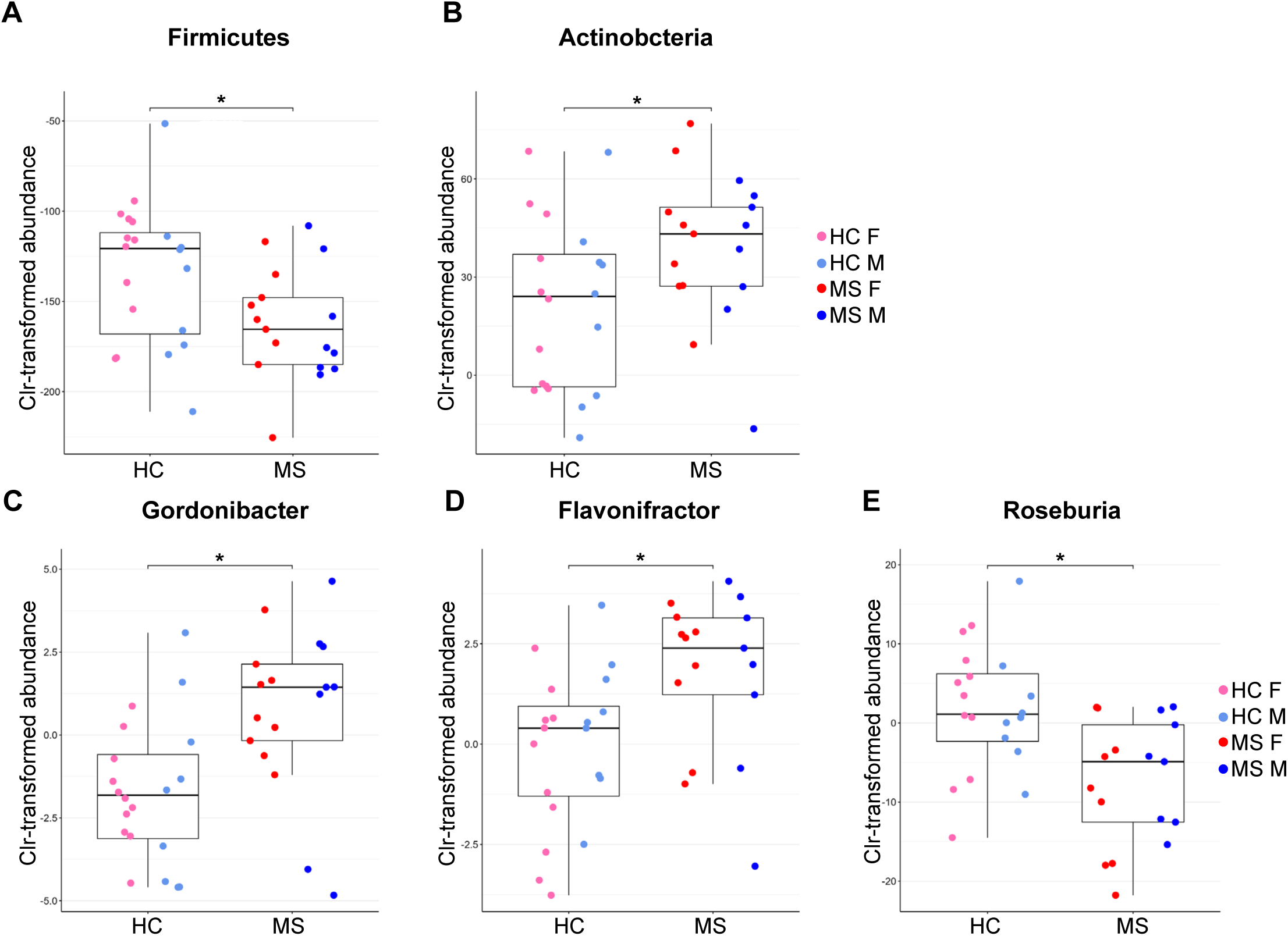
Microbial taxonomic profile of PwMS reveals a decrease in butyrate-producing bacteria. (A-E) Bar plot of Clr-transformed abundance of specific bacterial strains in HC and MS, median + IQR. Colored dots represent the sex component (pink = female HC; light blue = male HC; red = female MS; dark blue = male MS). Wilcoxon rank-sum test, p-value <0,05, * FDR <0,1. (A) Firmicutes; (B) Actinobacteria; (C) *Gordonibacter*; (D) *Flavonifractor*; (E) *Roseburia*. (A-E) HC n=20; MS n= 17.

At the *genus* level, PwMS had increased *Flavonifractor*^28^ (phylum Firmicutes) and *Gordonibacter*^29^ (phylum Actinobacteria), with an FDR of 0.064. *Flavonifractor* are flavonoid-degrading bacteria^30^. Slightly above the cut-out of the FDR (p-value 0.132), we also found an increase in *Eggerthella* (Actinobacteria), a known GABA producer^31^. Meanwhile, we found that PwMS have a statistically significant decrease in *Roseburia* (FDR 0.064), a butyrate-producing bacterial *genus*^32^, and a decreased trend slightly above the cut-off of the FDR, of *Lachnospira* (FDR 0.1008), *Faecalibacterium*^29^(FDR 0.132), *Eubacterium* (FDR 0.132), and *Coprococcus* (FDR 0.132), as illustrated in Fig. 2B and Table S2. All the *genera* found less abundant in PwMS belong to the phylum Firmicutes. *Roseburia, Faecalibacterium, Lachnospira, Eubacterium*, and *Coprococcus* are strongly linked to SCFAs synthesis, in particular butyrate^33^, strongly suggesting that the intestinal butyrate production is decreased in PwMS compared to HC. Furthermore, the same taxa have been found to correlate with MS also in other independent cohorts^3,34–36^, strengthening the biological significance of our data.

### Microbial functional analysis suggests an increase in glycolysis, stearate, and GABA synthesis in PwMS

Next, we explored the functional potential of the MS gut metagenome using the HUMAnN 3.9 workflow^37^ (Supplementary Table 3). No significant differences were observed after FDR correction, although two pathways were particularly increased in PwMS: super pathway of cytosolic glycolysis, pyruvate dehydrogenase, and TCA cycle (PWY-5464, FDR 0.38), and stearate biosynthesis I (PWY-5972, FDR 0.38).

The increased PWY-5464 could potentially be a consequence of decreased local butyrate concentration: without sufficient butyrate, colonocytes may rely more on glucose or pyruvate for mitochondrial oxidation. However, this shift is less efficient and may lead to lipid accumulation due to altered pyruvate flux^38^.

The stearate biosynthesis pathway refers to the metabolic process through which stearic acid, a long-chain saturated fatty acid (LCFA), is synthesized. Stearate is a key component of lipid metabolism and serves as a precursor for various biological molecules, including other fatty acids and complex lipids. It plays an essential role in cellular membrane structure and energy storage. Elevated concentrations of stearate have been associated with increased intestinal barrier permeability. Stearate-induced oxidative phosphorylation (OXPHOS) uncoupling, reduced ATP production, and elevated mitochondrial reactive oxygen species (ROS) lead to energy deprivation in enterocytes^39^.

MS pathogenesis has been closely associated with gastrointestinal dysfunction and leakiness of the intestinal barrier permeability^40^. By analyzing the microbiota taxonomical and functional profile, we found a decrease of SCFAs levels, particularly butyrate species, and increased levels of LCFA in PwMS compared to HC, suggesting a potential alteration of the intestinal barrier. To further investigate this aspect, we measured plasma intestinal fatty acid binding protein (FABP2) levels, a circulating biomarker of intestinal damage^41–43^. The results indicated no significant increase in intestinal permeability, as assessed by unpaired t-test or one-way ANOVA, at least at the disease stage we examined (Fig. S3).

We then conducted a more detailed analysis of the functional profile by examining human gut metabolic modules (GMM) based on Vieira-Silva et al.^44^ and the human gut-brain modules (GBM) based on Valles-Colomer et al.^45^ (Fig. 3). The GMM pipeline revealed a statistically significant increase in MF0031 – the glutamate degradation II pathway (Fig. 3A, B). Similarly, the GBM pipeline confirmed an increase in MGB022 – GABA synthesis III, although this did not pass FDR correction (FDR 0.2013; p-value 0.0096) (Fig. 3C, D). GABA is a key neurotransmitter that can be produced by bacteria expressing the glutamic acid decarboxylase (GAD, K01580) enzyme gene, responsible for converting glutamate to GABA. These bacteria belong primarily to the *phyla* Bacteroidetes and Actinobacteria^46^. Dysregulation in GABA-producing bacteria has been linked to several neurological and psychological disorders, although the mechanisms by which peripheral GABA influences brain function remain poorly understood. Since GABA is unlikely to cross the BBB^47^, peripherally produced GABA almost certainly influences brain function indirectly, primarily through the activation of GABA receptors in the enteric nervous system^48^.

**Figure 3.**
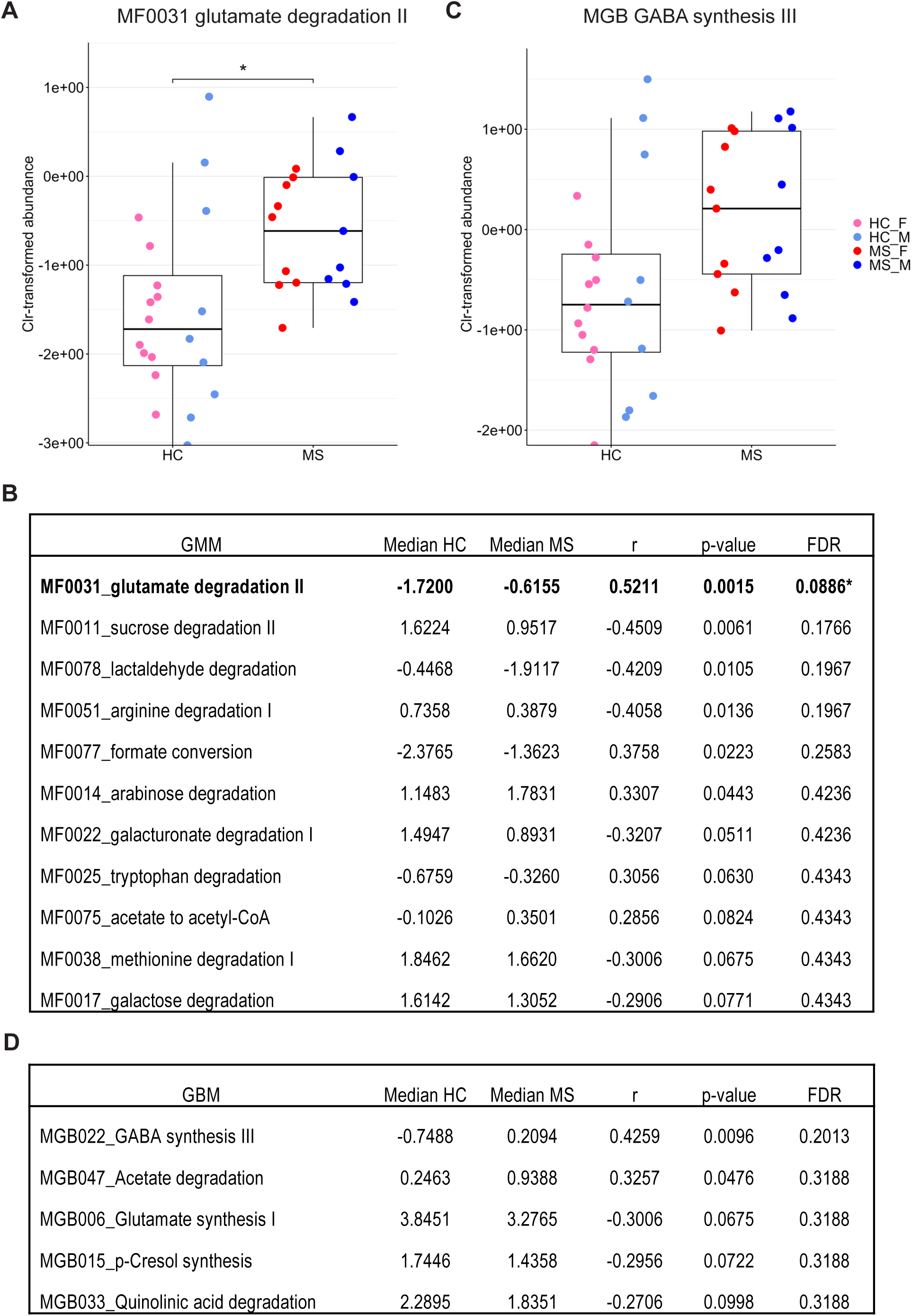
Functional analysis of stool microbiota reveals an increase in GABA synthesis in PwMS compared to HC. (A) Bar plot of Clr-transformed abundance of bacterial strains related to MF0031 pathway – glutamate degradation II - in HC and MS, median + IQR. Colored dots represent the sex component (pink = female HC; light blue = male HC; red = female MS; dark blue = male MS). (B) Statistics on GMM bacterial pathway analysis. r= effect size. Pathways in bold are statistically significant (* FDR <0,1). (C) Bar plot of Clr-transformed abundance of bacterial strains related to MGB022 pathway – GABA synthesis II - in HC and MS, median + IQR. Colored dots represent the sex component. Statistics on GBM bacterial pathway analysis. r= effect size. Pathways in bold have a p-value <0,05. (A-D) Wilcoxon rank-sum test, p-value <0,05, *FDR <0,1, HC n=20; MS n=17.

### The stool metabolomic profile partially reflects the microbial functional landscape

To further investigate the interaction of gut microbiota alterations and disease onset in PwMS, we performed metabolomic profiling in treatment-naïve PwMS and HCs. Untargeted metabolomics analysis was performed analyzing stool and urine samples using nuclear magnetic resonance (NMR). Stool samples did not reveal significant metabolomic differences between PwMS and HC, as shown by the score plot of PLS-DA considering both disease and sex differences (Fig. 4A, Q^2^= 0.16). Thus, the global metabolomic stool profile does not appear to be informative of the disease state (Fig. 4A).

**Figure 4.**
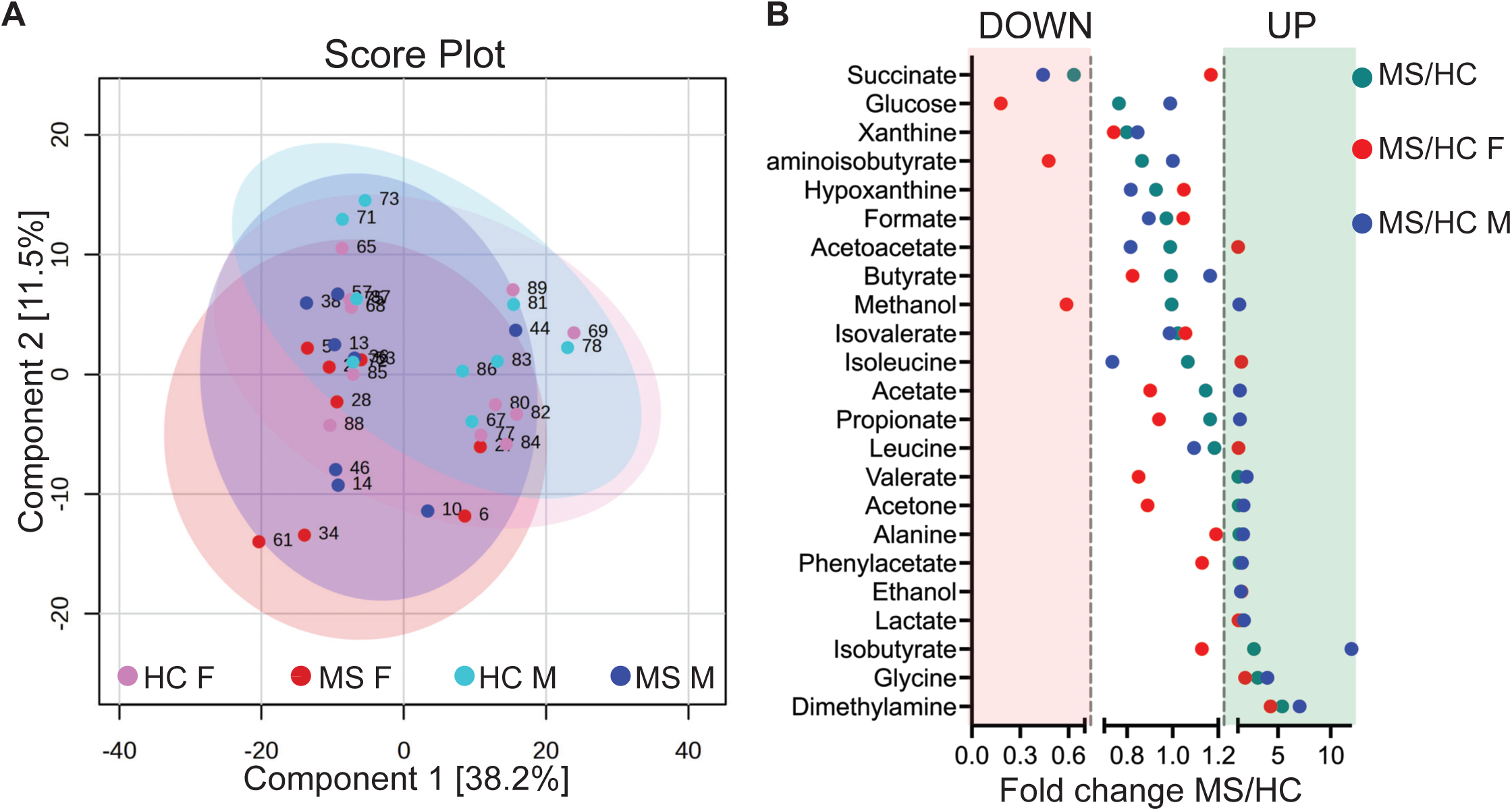
Stool metabolomic differences between PwMS and HC partially mirror microbial functional profiles. (A) PLS-DA of the stool untargeted metabolomic profile obtained by NMR on the four donor categories, represented by the colored dots, considering both disease and sex differences (pink = female HC; light blue = male HC; red = female MS; dark blue = male MS). Numbers represent the sample’s ID. Q^2^ (goodness of prediction) = 0.16. (B) Fold change (FC) of mean stool metabolite concentration measured by NMR in MS vs HC pooled (green dots), and sex-specific: male PwMS vs male HC (blue dots); female PwMS vs female HC (red dots). Downregulated metabolites: FC 0-0.7; upregulated metabolites: FC >1.2. (A-B) HC n=20; MS n= 18. HC F (female HC) n=11; HC M (male HC) n=9; MS F (female MS) n=9; MS M (male MS) n=9.

We subsequently assessed differences at the level of individual metabolites. Although no metabolite reached statistical significance following FDR correction (two-way ANOVA with Tukey correction or multiple Mann-Whitney test with Benjamini, Krieger, and Yekutieli correction), we noted patterns of potential interest regarding fold change (FC) of specific metabolites in MS versus HC (Fig. 4B). Succinate levels were decreased in PwMS, both when analyzing the overall disease group and when comparing only male donors. This finding aligns with the prediction from the functional microbiome profile derived from shotgun sequencing (Table S3), which showed increased succinate fermentation (PWY-5677) in PwMS. Conversely, PwMS showed increases in valerate, acetone, alanine, phenylacetate, ethanol, lactate, glycine, and dimethylamine. Considering the sex component, ethanol, lactate, glycine, and dimethylamine were consistently upregulated in PwMS (Fig. 4B), suggesting that these metabolites represent a disease-driven signature independent of sex. The increase in valerate, acetone, alanine, acetate, and phenylacetate, however, was exclusive to the male component of the cohort and absent in the female counterpart, suggesting a strong sex-dependent component (Fig. 4B). To our knowledge, none of these metabolites have previously been linked to MS pathogenesis. However, a previous study correlated increased levels of amino acids such as glutamine, arginine, and histidine in the stool of PwMS, suggesting a generally impaired amino acid utilization or microbial processing^49^. We also observed an increased fold change of isobutyrate in PwMS compared to HC. Isobutyrate is a branched SCFA that can be found in human stool as a result of the degradation of certain amino acids, particularly valine^50,51^. Interestingly, we found an increase in acetate only in male PwMS (two-way ANOVA with Tukey correction), which may reflect a potential shift in microbial fermentation, possibly related to changes in butyrate levels ^52,53^.

While we did not detect significant differences in butyrate or propionate levels themselves, certain metabolites, particularly lactate and phenylacetate, may indicate an altered SCFA-producing capacity. Lactate in stool is primarily produced by gut bacteria and serves as an important indicator of gut health and microbial composition. In healthy adults, lactate levels in the colon are typically low due to the presence of lactate-utilizing bacteria that rapidly convert lactate into propionate and butyrate^54^. The accumulation of lactate can lead to shifts in microbial community composition. A lower pH in the gut can exacerbate lactate accumulation, leading to decreased production of SCFAs and further shifts in microbial populations^55^. This situation can create a less stable microbial community, potentially contributing to inflammation and metabolic disorders^56^.

Phenylacetate is produced through the microbial degradation of unabsorbed phenylalanine in the gut^57^. Phenylacetic acid has been associated with a more pro-oxidant and immune-stimulated status, and it is negatively correlated with intestinal propionate^57^.

To conclude, although the stool metabolomic profile presents some features in line with the microbiome profiling (*i.e.,* succinate decrease in MS), it cannot be generally considered a good proxy for the intestinal metabolomic landscape. Indeed, the human stool metabolomic profile does not fully reflect the intestinal metabolomic profile due to significant spatial, compositional, and methodological differences^58^. Moreover, some metabolites like SCFAs are rapidly absorbed and metabolized or modified in the gut, meaning that stool only reflects residual or conjugated forms. Indeed, it is known that stool contains roughly only 5% of the total intestinal SCFAs production^59,60^. In conclusion, stool metabolomics offers practical insights but should not be equated with intestinal profiles due to inherent biological and technical limitations.

### Female PwMS present a unique urine metabolomic profile

To further investigate the link between disease onset, microbiome dysregulation, and metabolomic consequences at the systemic level, we performed untargeted metabolomics by NMR in urine samples. Contrary to stool samples, we found a global metabolomic profile alteration in PwMS compared to HC, as shown in the score plot of PLS-DA, when taking into account both disease and sex differences in the analysis (Q^2^= 0.48) (Fig.5A). Thus, we investigated which groups were driving the separation between PwMS and HC, by performing multiple PLS-DA slitted by the sex or disease component (Fig.5B-E): on MS donors discriminated by sex (Fig.5B), Q^2^= 0.05; on HC discriminated by sex (Fig.5C), Q^2^= 0.014; on male subjects discriminated by disease (Fig.5D), Q^2^<0; and on female donors discriminated by disease (Fig.5E), Q^2^= 0.54.

**Figure 5.**
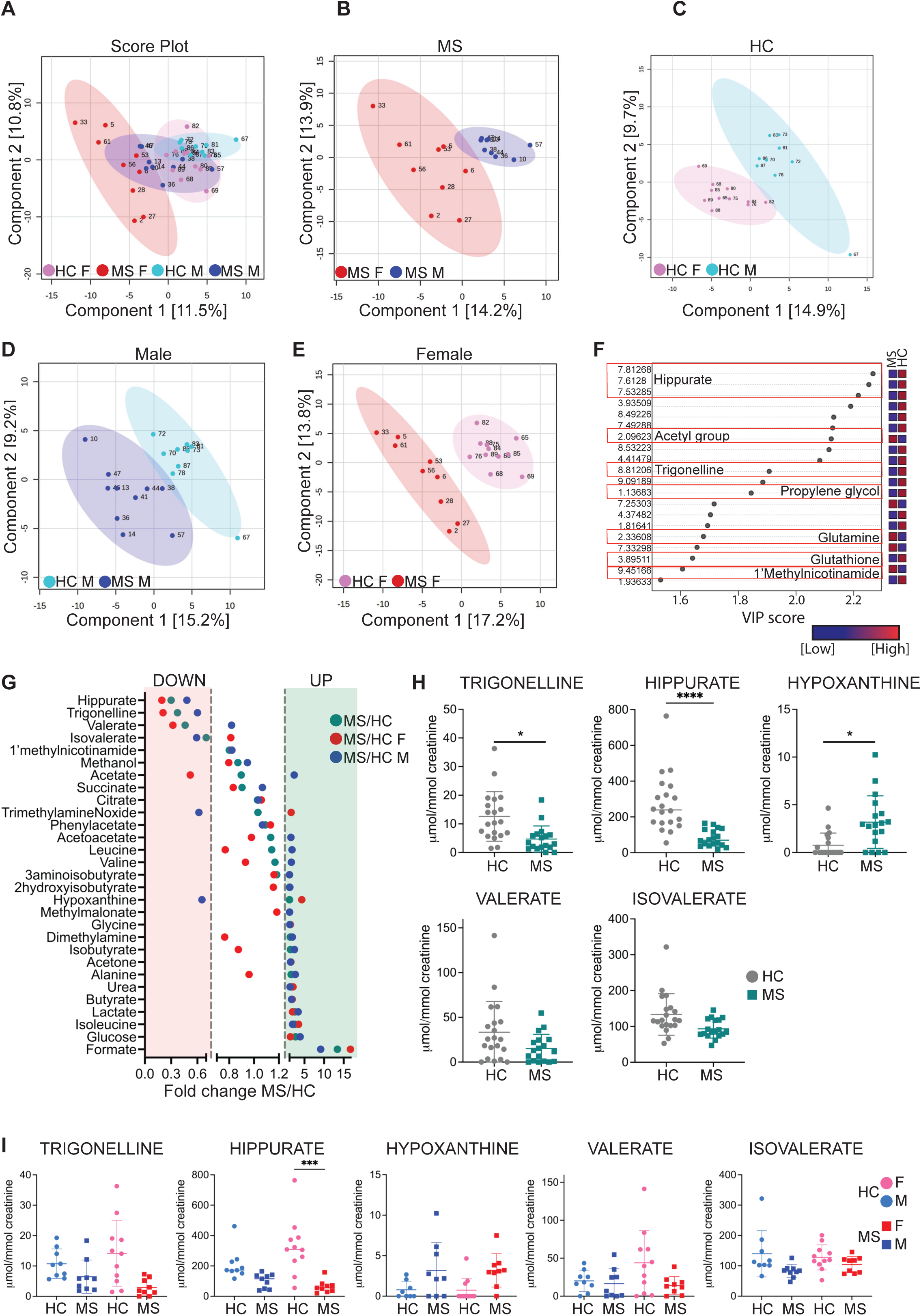
Female PwMS present a unique urine metabolomic profile. (A-E) PLS-DA of the urine untargeted metabolomic profile obtained by NMR, considering different donor categories. Colored dots represent the donor’s category (pink = female HC; light blue = male HC; red = female MS; dark blue = male MS), while numbers represent the sample’s ID. (A) PLS-DA on the four donor categories, considering both the disease and sex component; Q^2^= 0.48. (B) PLS-DA on MS donors discriminated by sex; Q^2^= 0.05. (C) PLS-DA on HC discriminated by sex; Q^2^= 0.014. (D) PLS-DA on male subjects discriminated by disease; Q^2^<0. (E) PLS-DA on female donors discriminated by disease; Q^2^= 0.54. (F) VIP scores of the female PLS-DA showing the metabolites that drive the separation between female HC and female MS. Numbers represent the bins related to the VIP scores. Bins corresponding to unidentified metabolites are also reported. The red (high) or blue (low) squares on the right indicate the level of expression in MS and HC. (G) Fold change (FC) of mean urine metabolite concentration measured by NMR in MS vs HC pooled (green dots), and sex-specific: male PwMS vs male HC (blue dots); female PwMS vs female HC (red dots). Downregulated metabolites: FC 0-0.7; upregulated metabolites: FC >1.2. (H-I) Urine metabolites of particular interest showing a different trend in MS and HC, considering only the disease discriminant (H), or disease and sex components (I), mean with SD. (H) Mann-Whitney test corrected for FDR with Benjamini, Krieger, and Yekutieli, *p < 0.05; ****p < 0.0001. (I) Statistical analysis with two-way ANOVA corrected for FDR with Tukey’s test; ***p < 0.0005. (A-I) HC n=20; MS n= 18. HC F (female HC) n=11; HC M (male HC) n=9; MS F (female MS) n=9; MS M (male MS) n=9.

Thus, the model can clearly distinguish between the groups and achieve predictive power only when considering all four groups together or female donors alone, implying that female MS and female HC carry the biggest metabolomic difference and drive the overall separation between MS and HC.

At this stage, we aimed to identify the specific metabolites driving the separation between female MS and female HC. The VIP scores identified hippurate, acetyl group, trigonelline, propylene glycol, glutamine, glutathione, and 1-methyl-nicotinamide as driving the groups’ uncoupling (Fig. 5F).

We then conducted a more detailed analysis at the single metabolite level to uncover subtle differences between PwMS and HC (Fig. 5G-I). Hippurate and trigonelline were confirmed to be significantly downregulated also by univariate analysis and FDR correction (multiple Mann-Whitney test coupled with Benjamini, Krieger, and Yekutieli correction), while hypoxanthine was increased in the urine of PwMS compared to HC (Fig. 5H). When accounting for sex, most metabolites showed similar trends in both sexes; however, fold changes were more pronounced in female donors (Fig. 5G, I), indicating a stronger disease-associated phenotype in women, consistent with findings from the metagenomic data. For instance, hippurate was statistically significant in female donors (PwMS versus HC) even after FDR correction, whereas no such significance was observed in the male cohort(two-way ANOVA with Tukey correction) (Fig. 5I).

Hippurate, a hepatic phase II conjugation product of microbial benzoate metabolism, has been associated with a healthy microbiota, particularly with microbial gene richness^61,62^. Thus, the decrease in urinary hippurate observed in PwMS compared to HC, especially in female donors, mirrors the trend of decreased alpha diversity in female PwMS compared to female HC, as identified by shotgun metagenomic analysis (Fig. 1A). Trigonelline, which cannot be produced by human cells, is synthesized through the methylation of nicotinic acid (niacin or vitamin B3) and acts as an NAD+ precursor, critical for cellular energy production and mitochondrial function. Decreased trigonelline levels in urine may reflect alterations in metabolic or physiological processes linked to muscle health^63^, NAD+ metabolism, and visceral fat dynamics. ^74^Trigonelline has been shown to inhibit the gut microbiota’s metabolism of choline, specifically reducing the production of trimethylamine (TMA) and its conversion to trimethylamine-N-oxide (TMAO)^64^. Indeed, we observe an increase in TMAO in female PwMS compared to female HC (Fig. 5G). Interestingly enough, trigonelline has been associated with neuroprotection^65–67^.

Hypoxanthine, which was elevated in MS compared to HC, is a purine base produced during the breakdown of adenine and guanine nucleotides, mainly generated during purine metabolism^68^. The increase in urine hypoxanthine aligns with the increase in PRPP-PWY and DENOVOPURINE2-PWY pathways found in the functional microbiome profile (Table S3): both of them are central to hypoxanthine synthesis, with the former supplying a critical precursor and the latter directing its conversion into hypoxanthine. Hypoxanthine levels rose in conditions of tissue hypoxia or oxidative stress, contributing to the generation of reactive oxygen species (ROS). It also serves as a marker of impaired ATP metabolism, where ATP is broken down into hypoxanthine and other purine degradation products. Overall, elevated urinary hypoxanthine may indicate tissue hypoxia, disorders of purine metabolism, or oxidative stress-related conditions.

Examining the fold changes of MS versus HC (Fig. 5G), a larger number of metabolites appeared differentially expressed, although they did not reach significance after FDR correction. In particular, PwMS show increased levels of glycine, glucose, acetone, butyrate, lactate, urea, isoleucine, and formate, representing a sex-independent, disease-driven metabolic signature.

Few studies have examined the urine metabolomic profile of PwMS compared to HC^69,70^. One study comparing HC, PwMS, and neuromyelitis optica also identified distinct metabolic signatures in the urine of PwMS. Interestingly, most metabolites that constitute the MS-driven signature overlap with our findings, but exhibit the opposite trend, likely because their cohort included previously diagnosed patients already under DMTs. As previously reported by the iMSMS Consortium^3^, DMTs have a significant impact on both the microbiota and the systemic metabolomic profile^69^.

In conclusion, we found that the urine metabolomic profile is strongly influenced by the microbiome’s taxonomic and functional composition, displaying MS-specific features, with females showing a more pronounced phenotype than males.

### PwMS have increased plasma concentration of total SCFAs, and altered SCFAs relative ratio

Having established that the microbiome of PwMS differs significantly from that of HC, with measurable effects on the circulating metabolome, we next investigated whether we could further characterize changes in SCFAs and other microbiome-related metabolites, particularly those implicated in MS-associated taxonomic alterations and previously linked to MS onset and pathogenesis. We thus extracted SCFAs values from stool and urine NMR profiles, and specifically measured SCFAs concentration in the plasma and the cerebrospinal fluid (CSF) by gas chromatography-mass spectrometry (GC-MS). For ethical reasons, CSF sampling was limited to PwMS. First, we calculated the amount of total SCFAs, given by the sum of the concentration of the single SCFAs found in each specific biofluid, and compared PwMS with HC, considering both the overall disease group (Fig. 6A) and also the sex-component (Fig. 6B). We found that PwMS had a significant increase in total plasma SCFAs (unpaired t-test). Both male and female donors showed an increased trend, indicating this is a general, sex-independent, disease-driven feature, although statistical significance was not reached due to limited sample size (one-way ANOVA; F MS vs F HC p = 0.1; M MS vs M HC p = 0.18) (Fig. 6B).

**Figure 6.**
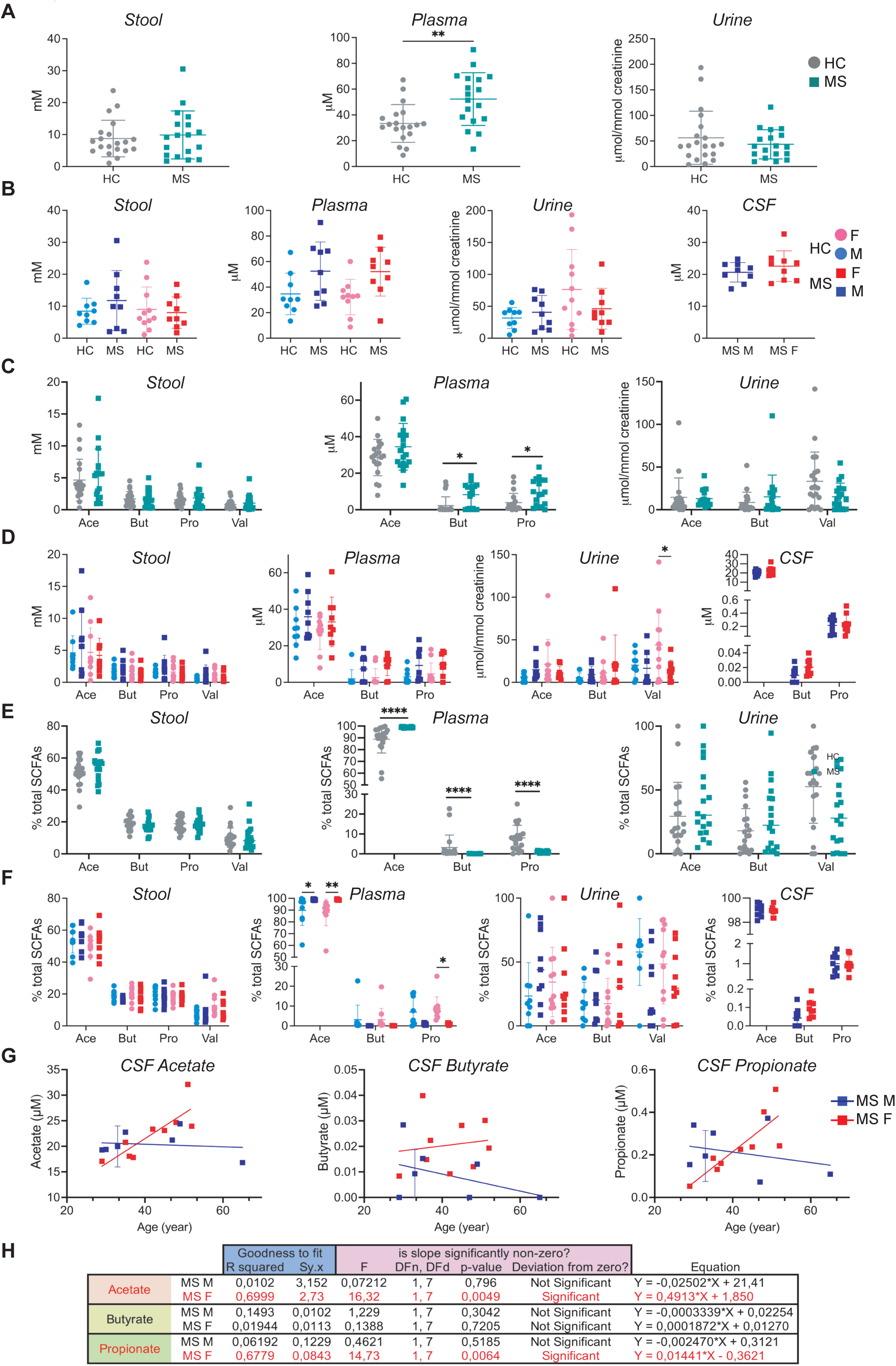
District-, sex-, and age-specific SCFAs alterations in PwMS. (A-B) Concentration of total SCFAs in stool, plasma, and accounting only for the disease state (A) or also for the sex component (B). Here, we also added the total SCFAs concentration in the CSF. (C-D) Concentration of every single metabolite in stool, plasma, and urine, accounting only for the disease state (C) or also for the sex component (D). Here, we also added the single metabolite concentration in the CSF. (E-F) Percentage of acetate (Ace), butyrate (But), propionate (Pro), and valerate (Val) on total SCFAs concentration, accounting only for the disease state (E) or also for the sex component (F). Here, we also added the percentage of acetate, butyrate, and propionate of the total SCFAs concentration in the CSF. (A-F) mean with SD. (A) unpaired t-test; (B) one-way ANOVA; (C-F) statistical analysis between two groups: Mann-Whitney test corrected for FDR with Benjamini, Krieger, and Yekutieli correction; statistical analysis between four groups: two-way ANOVA corrected for FDR with Tukey. *p < 0.05, **p < 0.01, ****p < 0.0001. (G-H) Sex-specific correlations between CSF acetate, butyrate, and propionate and the donor’s age, tested by a simple regression model. (G) Correlation plot with interpolation line and (H) regression model calculation. Significant values are in bold. r^2^= coefficient of determination; Sy.x= standard error of regression; F= F score; DFn,DFd= degrees of freedom numerator, degrees of freedom denominator. (A-H) HC n=20; MS n= 18. HC F (female HC) n=11; HC M (male HC) n=9; MS F (female MS) n=9; MS M (male MS) n=9.

We then analyzed each metabolite in detail to understand how individual concentrations were altered in PwMS compared with HC (Fig. 6C-D), finding a significant increase in butyrate and propionate in the plasma of PwMS, while the increase of acetate levels did not reach statistical significance after FDR correction (Mann-Whitney test with Benjamini, Krieger, and Yekutieli correction) (Fig. 6C). As seen previously with total SCFA concentration, the trend was shared between male and female donors, but statistical significance was not achieved when considering sex (Fig. 6D; two-way ANOVA with Tukey correction).

We then investigated whether both the absolute concentrations and the physiological ratios of SCFAs were altered in PwMS. Each anatomical district presents a precise ratio that varies from 60:20:20 (acetate: propionate: butyrate) in the intestine, to 90:5:5 in the systemic circulation^71,72^. We calculated the percentage of each metabolite relative to the total SCFA amount and found that, although the overall amount of butyrate and propionate increased, their percentage decreased in PwMS compared to HC, while the percentage of acetate increased (Figure 6E, Mann-Whitney test coupled with Benjamini, Krieger, and Yekutieli correction). In our cohort, we obtained a plasma acetate: propionate: butyrate ratio of 88.8±11.7:7.9±6.3:3.1±6.3 for HC, and a ratio of 98.9±0.56:0.99±0.43:0.06±0.05 for PwMS. When the sex component was considered, we confirmed the increased percentage of acetate in PwMS of both sexes and the decrease in propionate in females (Figure 6F, two-way ANOVA coupled with Tukey correction). Decreased levels of butyrate in both sexes, and of propionate in male PwMS did not reach statistical significance (Fig. 6F, two-way ANOVA with Tukey correction).

### Distinct SCFA alterations in PwMS across region, sex, and age

Next, we explored sex- and district-specific differences in SCFA bioavailability. While the core SCFA signature was consistent across sexes, certain metabolites exhibited sex-dependent variations. In females, butyrate increases were more pronounced in both plasma and urine, whereas in males, the rise was primarily observed in plasma. Analysis across anatomical districts revealed that these alterations were not uniform: some regions showed stronger elevations in specific SCFAs, highlighting a complex interplay between anatomical site, sex, and disease status (Fig. S5A–D). Male PwMS presented an overall upregulation in SCFAs in all the districts considered, except for stool butyrate and urine valerate FC, which remained unvaried (Fig. S4A).

Regarding the CSF, we compared individual SCFAs between male and female PwMS and observed higher butyrate concentrations in female donors. Although the underlying biological mechanism remains unclear, it is noteworthy that, despite comparable plasma butyrate levels in males and females, female PwMS exhibit elevated CSF butyrate. This finding suggests that CNS butyrate metabolism may be influenced by sex-dependent regulatory factors and warrants further investigation

Because both microbiota and microbiota-related metabolites can be influenced by sex and age, we examined whether donor age affected group identity (MS or HC), sex, or individual SCFA concentrations in each biofluid. Although our metagenomic analysis did not reveal any association between age and microbiome profiles, we assessed age correlations with SCFA levels. No significant correlations were observed in stool (Fig. S5A, B), plasma (Fig. S5C, D), or urine (Fig. S5E, F) using a simple regression model, indicating that age is not a confounding factor for these biofluids. Interestingly, in CSF, acetate and propionate concentrations increased with age in females but not in males (Fig. 6G, H), further supporting the notion that SCFA metabolism in the CNS may be modulated by sex-dependent factors

## DISCUSSION

Although it is well established that the gut microbiome is altered in PwMS compared with HC, our study provides new insights into the role of the microbiota-gut-brain axis in neuroinflammation, highlighting several distinguishing features. Importantly, we focused exclusively on a cohort of treatment-naïve, newly diagnosed PwMS, allowing us to capture microbiome and metabolomic alterations at disease onset while minimizing the confounding effects of disease-modifying therapies (DMTs) and accounting for sex differences. In addition, we performed comprehensive metagenomic profiling, examining not only the bacterial microbiome but also the virome and unicellular eukaryotes. Finally, by integrating metagenomic data with multi-fluid metabolomic analyses, we were able to trace the systemic consequences of microbiota dysbiosis across the entire body

As a general consideration, it is important to note that participants in our cohort were not following a standardized diet, but instead adhered to their habitual, free-choice dietary regimens. Since diet is a major determinant of microbiota composition, the presence of mostly omnivores, along with a few individuals on special diets (e.g., vegan or lactose-free), may partly account for intra-group variability at the taxonomic level, which also impacted the functional profile. Nevertheless, we were able to identify a core disease-driven microbiome signature that remained more consistent and stable than the variations attributable to dietary differences. Additionally, our findings underscore the importance of considering sex as a biological variable in human studies. Several features emerged as sex-specific or sex-modulated, indicating that pooling data from both sexes without prior stratification may obscure meaningful biological differences or lead to misinterpretation of results.

Concerning the metagenomic data, our analysis encompassed not only the bacterial component but also the virome and the protozoan Blastocystis. Regarding the bacterial microbiota, we did not observe any significant difference in alpha diversity between MS and HC groups as measured by different indexes (Observed, Simpson, Fisher, and Pielou) (Fig. 1A, B). However, a trend of decreased alpha diversity was observed in PwMS, predominantly females, compared to female HC. The small size of the cohort limited the statistical significance of our analysis; however, the data were supported by the reduced hippurate levels in the urine of female PwMS compared to female HC. Interestingly, the beta diversity analysis underlined that the group identity (HC or MS) and the disease status (“relapse”, “remission”, “progression”, and “healthy control”) are the only variables statistically correlating with microbiota variability (Fig. 1D).

At the taxonomic level, our study confirmed MS-associated microbiota alterations previously reported in the literature. Actinobacteria, found to be increased in PwMS, are known to influence the gut-brain axis through GABA^73^, as highlighted by the functional metagenomic analysis of GMM and GBM (Fig. 3). On the other hand, PwMS showed a decrease in Firmicutes (Fig. 2A), known producers of SCFAs, especially butyrate and propionate^74,75^. A decrease in intestinal butyrate production has profound consequences for the overall microbiota and intestinal homeostasis^76^ as butyrate serves as the primary energy source for colonocytes, providing 70–80% of colonocyte energy needs^77^. This energy deficit activates stress response signals in colonocytes^77,78^. Butyrate also helps maintain a hypoxic environment that supports the growth of anaerobic microbiota. Reduced butyrate levels decrease oxygen consumption, disrupting luminal anaerobiosis and potentially altering the composition of the gut microbiota ^78^. Moreover, butyrate helps stabilize the gut barrier by activating specific receptors (e.g., GPR41, GPR109A) and inhibiting histone deacetylases (HDACs)^79–81^. A decrease in butyrate compromises these protective mechanisms, increasing susceptibility to local inflammation^82–84^. Reduced butyrate levels can also trigger a negative feedback loop that limits local SCFA availability by impairing colonocyte uptake, through decreased expression of transporters such as monocarboxylate transporter 1 (MCT1) and sodium-coupled monocarboxylate transporter 1 (SMCT1)^85^. Consequently, lower intestinal butyrate concentrations promote the development of a dysbiotic environment^86^.

In our study, we also examined the virome component of the gut ecosystem. Although virome alpha diversity did not differ significantly between groups, we observed a trend toward increased diversity in PwMS compared with HC (Fig. 1E). This pattern mirrors findings from a previous study analyzing the virome in autoimmune disorders, which reported an overall reduction in viral diversity in systemic lupus erythematosus and rheumatoid arthritis but noted a trend toward increased variability specifically in MS^87^. Conversely, another published analysis described the opposite trend, although that cohort included patients undergoing DMT, a major confounding variable likely accounting for the discrepancy with our treatment-naïve population^88^.

Beyond the virome, we also assessed the presence of the gut protozoan Blastocystis. We detected four subtypes (1, 3, 6, and 9), exclusively in a small fraction of HC individuals and without any association with sex or age (Fig. S2C). These observations are consistent with recent work linking Blastocystis to a healthier gut microbial ecosystem. To our knowledge, this is the first study to investigate Blastocystis prevalence in the metagenome of PwMS, providing an additional layer of insight into non-bacterial components of the gut microbiome and their potential relevance to MS.

The metabolomic consequences of an altered microbiome were only partially reflected in the stool metabolomic profile, largely because most metabolites are rapidly absorbed, metabolized, or transformed within enterocytes. Conversely, the urine metabolomic profile, often overlooked in microbiota studies, proved to be highly informative. We observed an overall alteration in the metabolomic profile in PwMS compared to HC, as shown in the PLS-DA analysis. This was evident both when considering all four donor categories, accounting for disease status and sex (Q^2^= 0.48, Fig.5A), and when focusing exclusively on female donors stratified by disease status (Q^2^= 0.54, Fig.5E). These findings indicated that the most profound metabolomic differences occur between female PwMS and HC, suggesting that this subgroup largely drives the overall separation observed between MS and HC. Through the VIP score analysis, we observed that the separation between PwMS and HC was led by hippurate, a urinary marker of microbial alpha diversity, together with acetyl group, trigonelline, propylene glycol, glutamine, glutathione, and 1’methylnicotinamide (Fig. 5F). Systemic trigonelline has been associated not only to muscle strength but also to inflammation and oxidative stress, as it modulates Nrf2 and NF-κB pathways, thereby balancing oxidative stress and inflammatory responses. Thus, while not directly linked to urinary levels, systemic trigonelline deficiency might diminish its neuroprotective effects, likely reducing neuroinflammation and enhancing neurotransmitter levels (*e.g*., dopamine and serotonin). Previous studies analyzed urine metabolomics of PwMS with HC, although they recruited mostly patients undergoing DMTs^69,89^. Interestingly, prior comparisons across HC, PwMS, and patients with neuromyelitis optica revealed a distinct urine metabolomic profile in PwMS ^69^. Although this profile shares several features with our findings, it exhibits an opposite trend, most likely because that cohort consisted of previously diagnosed individuals who were already receiving treatment, a factor that may also explain earlier discrepancies noted for virome alpha diversity^69^. As was previously reported, DMTs have a strong impact on microbiota and systemic metabolomic profile^3^. These discrepancies reinforce the need to recruit cohorts that are carefully matched to the study’s scientific aims. Because our investigation targeted the disease-onset phase, before long-term progression or exposure to DMTs, it is expected that our results differ from those obtained in studies involving treated or more advanced cases.

Our analysis focused on SCFAs, key microbial metabolites that have previously been associated with MS. Although the stool metabolomic profile did not reveal lower levels of SCFAs in PwMS (Figure 6C, D), the shotgun metagenomic analyses predicted a decrease in major SCFA-producing bacteria such as Firmicutes (Table S2), suggesting a likely drop in SCFAs production, particularly butyrate. Interestingly, we found an opposite tendency. Notably, an opposite trend was observed in plasma, where propionate and butyrate levels were increased in PwMS; acetate followed the same trend, although it did not reach statistical significance. The total plasma SCFA concentration was also elevated. Importantly, dysregulation of individual plasma SCFAs was also observed (Figures 6E, F). As noted earlier, stool contains only roughly 5% of the total intestinal SCFA production and does not accurately reflect intestinal SCFAs concentration, as these metabolites are rapidly metabolized and absorbed by epithelial cells before entering the bloodstream^90^. Increased circulating SCFA levels, if not supported by higher intestinal synthesis, could result from either intestinal barrier leakage or impaired intestinal absorption. We ruled out the first hypothesis by showing that a circulating marker of intestinal leakage, FABP2, is expressed at comparable levels in both PwMS and HC (Fig. S3). Thus, we propose that decreased intestinal SCFAs production leads to a defect in intestinal SCFAs absorption and metabolism in the gut, thus increasing SCFAs bioavailability in the systemic circulation.

Circulating SCFAs are essential regulators of energy metabolism, barrier function, and immune homeostasis. Thus, disturbances in their systemic levels and ratios may have downstream effects on the immunometabolism of peripheral immune cells, which is compromised in PwMS.. Indeed, most of the studies agree on impaired glycolysis and mitochondrial respiration in T cells of PwMS^91–95^. Furthermore, circulating immune cells of PwMS show impaired mitochondrial redox status and deficient antioxidant capacity, suggesting an increase in circulating ROS^96,97^. Concomitantly, SCFAs have been shown to modulate immunometabolism by fueling the TCA cycle and oxidative phosphorylation (OXPHOS), both in T cells and in macrophages.

SCFAs in the CSF originate directly from the bloodstream, as these metabolites can cross the blood–brain barrier (BBB) via monocarboxylate transporters. Once within the CNS, SCFAs influence brain function through several mechanisms: they support BBB integrity by upregulating tight-junction proteins, modulate microglial activation with potential anti-inflammatory effects, and contribute to gut–brain axis communication by regulating neurogenesis, serotonin biosynthesis, and neuronal homeostasis. At a cellular level, SCFAs reduce ROS production under hypoxic conditions, stabilize mitochondrial membrane potential, and suppress caspase activation in neuronal cells. Serum SCFA levels also correlate with hippocampal brain-derived neurotrophic factor (BDNF), a key mediator of synaptic plasticity and cognitive function. Conversely, elevated systemic SCFA concentrations can alter phosphatidylethanolamine plasmalogens and cardiolipin metabolism, lipid classes essential for mitochondrial function, potentially exacerbating neurodegenerative processes. Thus, SCFAs in the CSF could act as double-edged swords: while they enhance resilience to acute injury and modulate neuroinflammatory processes, their dysregulation could contribute to neuronal dysfunction, underscoring the need for precision when considering SCFA-based therapeutic strategies. Notably, alterations in CSF SCFA levels have been reported across diverse neurological conditions, supporting their potential as biomarkers or therapeutic targets in disorders such as autism spectrum disorder and Alzheimer’s disease.

It is therefore important to recognize that, although SCFAs are generally associated with beneficial effects across various disorders, their impact is highly dependent on concentration, metabolic context, and anatomical location. Beyond their direct actions, SCFAs can also modulate the gut-brain axis indirectly by stimulating the enteric nervous system. While the precise molecular mechanisms remain to be fully elucidated, a reduction of intestinal SCFAs may compromise brain homeostasis by attenuating signaling through the enteric nervous system and the vagus nerve.

In summary, this study demonstrates that PwMS exhibit not only a distinct gut microbiota composition but also a unique urinary metabolomic profile compared with HCs. Notably, female PwMS show a pronounced reduction in urinary hippurate levels, reflecting decreased microbial alpha diversity. By employing an intra-donor, multi-fluid metabolomic approach, we traced the bioavailability of SCFAs across biological compartments, revealing a link between neuroinflammation and altered SCFA production, intestinal absorption, and systemic distribution. While these alterations appear to be primarily disease-driven, female donors display a more marked microbiome-related phenotype across multiple readouts. Overall, our work provides an in-depth characterization of how microbiome alterations impact the systemic metabolomic landscape in treatment-naïve PwMS at disease onset.

### Limitations of the study

We acknowledge several limitations of this study. With respect to the cohort, this was a monocentric study that did not account for geographical variability, and the number of enrolled subjects was relatively small (Table 1). The limited sample size represented a key constraint for achieving statistical significance in the metagenomic analyses; nonetheless, we were still able to identify core disease-associated features, supporting the robustness of our findings. In addition, BMI data were not available for all participants (missing BMI values for 5 HC).

As for the metabolomic component, our study lacks CSF metabolomic profiling in HC. Although such data would have been highly informative, collecting CSF from healthy individuals was not ethically feasible, as lumbar puncture is an invasive procedure performed exclusively for clinical indications. We were also constrained by technical considerations: obtaining a fully comprehensive metabolomic landscape would have required combining NMR, GC-MS, and LC-MS analyses on the same samples.

Conceptually, our study provides correlational evidence that, in newly diagnosed treatment-naïve PwMS, microbially derived dysregulation leads to systemic metabolomic modifications and provides testable targets for modifying the microbiome and its metabolites within the context of MS. Nonetheless, as a patient-based study, our work does not provide the mechanistic insights needed to directly link the dysregulated metabolomic profile to disease onset and pathogenesis. Future *in vivo* animal studies and human *in vitro* models will be essential to address this molecular gap.

## RESOURCE AVAILABILITY

### Lead contact

Further information and requests for resources and reagents should be directed to and will be fulfilled by the lead contact, Paola Panina-Bordignon.

### Materials availability

This study did not generate new, unique reagents.

### Data and code availability

- The datasets generated during the current study are not yet publicly available but will be deposited in GEO upon acceptance.
- Any additional information required to reanalyze the data reported in this paper is available from the lead contact upon request.

## ACKNOWLEDGMENTS

We thank Prof. Massimo Filippi, Director of the Neurology Unit, IRCCS San Raffaele Hospital, for granting access to patient samples; Centro Risorse Biologiche of the San Raffaele Hospital for handling human samples and storage, and for ensuring donor anonymization; Elena Brambilla and Carolina Peri for their technical assistance. This work was supported by a Marie Skłodowska-Curie Actions Postdoctoral Fellowship from the European Union (grant 101031375 to J.P.) and by the Fondazione Italiana Sclerosi Multipla (FISM) (grant RI0294-FISM-2022 to P.P-B.).

## AUTHOR CONTRIBUTIONS

J.P. conceived and designed the study, analyzed and interpreted the data, wrote the manuscript, secured funding, and supervised the overall project; Va.M. performed NMR experiments, analyzed and interpreted NMR data, and edited the manuscript; L.M., M.V-C., F.A., E.P. and A.G-V. processed and analyzed shotgun metagenomic data, interpreted results, and edited the manuscript; M.B. and D.G. performed GC-MS experiments and analyzed data; F.M., Vi.M. and M.F. recruited the donors, collected the biological samples and gathered all patient-associated clinical data; N.S. supervised the shotgun metagenomic sequencing and contributed to the manuscript editing; G.M. contributed to manuscript editing and critical revision; P.P-B. supervised the overall project and data analysis, wrote and edited the manuscript, and secured funding.

## DECLARATION OF GENERATIVE AI AND AI-ASSISTED TECHNOLOGIES

During the preparation of this work, the authors used Perplexity.ai to assist with text editing. All content generated using this tool was subsequently reviewed and revised by the authors, who take full responsibility for the content of the publication.

## KEY RESOURCES TABLE

| REAGENT OR RESOURCE | SOURCE | IDENTIFIER |
| --- | --- | --- |
| Biological samples |  |  |
| Human healthy controls and MS subjects' stool | This paper | CRB biorepository, San Raffaele Hospital |
| Human healthy controls and MS subjects' urine | This paper | CRB biorepository, San Raffaele Hospital |
| Human healthy controls and MS subjects' plasma | This paper | CRB biorepository, San Raffaele Hospital |
| Human MS subjects' cerebrospinal fluid | This paper | CRB biorepository, San Raffaele Hospital |
| Critical commercial assays |  |  |
| DNeasy PowerSoil Pro Kit | Qiagen | Cat # 47014 |
| Illumina® DNA Prep, (M) Tagmentation kit | Illumina | Cat # 20060059 |
| Metabolomics on stool and urine | This paper | Center for Omics Sciences, San Raffaele Hospital |
| SCFAs analysis on plasma and cerebrospinal fluid | This paper | Institute of Bioimaging and Complex Biological Systems, National Research Council |
| Software and algorithms |  |  |
| Prism 10 | GraphPad | version 10.4.2 |
| BioRender | BioRender | https://biorender.com/n08n589 |
| MestreNova 14.1.2 | Mestrelab Research S.L |  |
| Chenomx 8.6 | (Chenomx, Inc., Edmonton, AB, Canada) |  |

## EXPERIMENTAL MODEL AND STUDY PARTICIPANT DETAILS

A total of 38 PwMS and HCs were included in this study. See Table 1, Table S1, and Figure S1 for phenotypes of all participants. Participants were all recruited at San Raffaele Hospital (Milan, Italy). Inclusion criteria for PwMS required that participants carry a diagnosis of MS or CIS (McDonald et al., 2001^98^), be newly diagnosed, and still treatment naïve. HCs are genetically unrelated to PwMS; they were recruited to match as much as possible the sex and age distribution of PwMS. Exclusion criteria for PwMS and HC included the presence of other autoimmune disorders, diagnosed gastrointestinal or neurological disorders. Participants were excluded if they received antibiotics or corticosteroids within the past three months before sample collection. Participants were not on a standardized diet but were following a free regimen. Human subject research approval was obtained through the San Raffaele Hospital’s ethics review committee, protocol BRAINESS, n. 131/INT/2021. All participants provided written informed consent before sample collection. All research involving human participants has been planned, conducted, and reported following the WMA’s Declaration of Helsinki as revised in 2024.

## METHOD DETAILS

### SPECIMEN COLLECTION

Samples from PwMS (urine, stool, blood) were all collected by the nursing staff at the Neurology Department at San Raffaele Hospital and rapidly sent to the Biobank Center (CRB). The CSF was obtained from the leftover of the clinical exam and collected within 3 days of the other biofluids. Concerning HC, blood was collected by specialized nursing staff at San Raffaele Hospital, while urine and stool were collected by the donor at home through a collection kit. Urine and stool were collected early in the morning, the same day as blood withdrawal, and kept and transported at 4°C for a maximum of 4h before processing. All samples were collected, anonymized, and stored at the CRB. CSF samples are collected from each subject by rachid puncture and collected into a sterile polypropylene tube. The samples were centrifuged at 1200 rpm for 10 minutes at room temperature (RT) to separate the cellular component. The supernatant was then aliquoted, snap-frozen in liquid nitrogen, and stored at −80 °C. Blood for plasma preparation was collected in tubes containing lithium heparin. Samples were centrifuged at 1,000g for 10 minutes at 4°C. The supernatant was aliquoted and stored at −80°C. Urines were centrifuged at 1600 g for 10 min at 4 °C to remove any possible cellular contamination, then aliquoted and stored at −80°C. Stools were aliquoted and stored at −80°C. PwMS were required to complete a clinical survey to report the disease status and clinical parameters (Table S1), and all participants completed the subject survey to report medication, lifestyle, and pre-existing extra-intestinal pathologies (Table 1). Clinical outcomes included the EDSS^99^. All data were collected and stored anonymously through the donor’s unique ID code.

### WHOLE METAGENOME SHOTGUN SEQUENCING

DNA was extracted from aliquots of fecal samples using the DNeasy PowerSoil Pro Kit (Qiagen) following the manufacturer’s instructions. Sequencing libraries were prepared using the Illumina® DNA Prep (M) Tagmentation kit (Illumina), following the manufacturer’s guidelines. Sequencing was performed on a NovaSeq6000 S4 flow cell (Illumina) as a paired-end 150 nucleotide-cycle run targeting a sequencing depth of 7.5 Gbp per sample at the internal sequencing facility at the University of Trento, Trento, Italy.

### METAGENOMIC DATA PRE-PROCESSING AND QUALITY CONTROL

Reads from stool samples were pre-processed using the pipeline described in https://github.com/SegataLab/preprocessing. Shortly, metagenomic reads were quality controlled and reads of low quality (quality score <Q20), fragmented short reads (<75 bp), and reads with >2 ambiguous nucleotides were removed with Trim Galore (v0.6.6). Contaminant and host DNA were identified with Bowtie2 (v2.3.4.3)^100^ using the -sensitive-local parameter, allowing confident removal of the Instrument’s spike-ins and host-associated reads (hg19 human genome release). Only samples with more than 30 million reads after the preprocessing were included in the downstream analysis (N=37, median=62.5 million reads/sample, IQR=21.5 million reads/sample). Read statistics of all samples (number of reads, number of bases, minimum, median, and maximum read length per sample are detailed in Table S4).

### TAXONOMIC AND FUNCTIONAL MICROBIOME PROFILING

Species-level profiling was performed on all the samples with MetaPhlAn 4.1.1^101^ with default parameters and the custom database (available at http://cmprod1.cibio.unitn.it/biobakery4/metaphlan_databases/). Functional profiling was performed with HUMAnN 3.9^37^, using default parameters and the Jun23 CHOCOPhlAnSGB_202403 database. MetaCyc^102^ pathway definitions were reported as provided by HUMAnN. Only features with an average relative abundance of at least 0.0001 were kept. Gut-brain module (v1.0)^103^ and gut metabolic module (v1.07)^44^ abundances were calculated with omixer-rpmR (v0.3.3)^104^. Only modules with an average relative abundance of at least 0.001 were kept.

### ALPHA DIVERSITY, BETA DIVERSITY, AND ORDINATION

Species-level abundance matrices obtained in MetaPhlAn 4.1.1 were centered log ratio-transformed using the codaSeq.clr function in the CoDaSeq R package (v0.99.7)^105^, using the minimum proportional abundance detected for each species for the imputation of zeros. Only species with an average relative abundance of at least 0.0001 were kept. Species-level richness (Observed) and diversity (Simpson’s) were determined using the estimate_richness function in phyloseq, and evenness (Pielou) was calculated as Shannon’s diversity divided by the logarithm of the Observed richness^106^. A PCoA on Aitchison distance (Euclidean distance between samples after centred log-ratio transformation) was produced with the ordinate and plot_ordination functions in phyloseq (v1.50.0)^107^. The association between metadata variables and distance matrices was assessed by PERMANOVA with the capscale function in vegan (v2.6.10)^108^.

### BLASTOCYSTIS PROFILING

The presence of Blastocystis in metagenomic samples was assessed using a previously validated computational workflow^109^ as in Piperni E. et al^19^. Nine reference genomes for eight different Blastocystis subtypes (i.e., ST1 [ST1_LXWW01], ST2 [ST2_JZRJ01], ST3 [ST3_JZRK01], ST4 [GCF_000743755 & ST4_BT1_JZRL01], ST6 [ST6_JZRM01], ST8 [ST8_JZRN01], and ST9 [ST9_JZRO01]) were mapped against metagenomic reads with Bowtie2 (v2.3). SAMtools (v1.19) and bedtools were then used to compute the breadth of coverage of each genome, and a sample is considered to be positive for a Blastocystis subtype if the respective genome has a breadth of coverage of at least 10%.

### VIROME PROFILING

The presence of bacteriophages in metagenomic samples was assessed using MetaPhlAn 4.1.1^110^ with the parameter--profile_vsc. Then, the presence/absence of viruses was assessed between MS patients and healthy controls with the Fisher test, followed by multiple testing corrections.

### URINE AND STOOL NMR METABOLOMICS

Urine samples were thawed at room temperature for 30 minutes. 900μl of urine was mixed with 100μl of buffer (1.5M KH2PO4/D2O, pH 7.4 (with KOD/D2O), 2mM NaN3, 0.1% TSP (3-(trimethyl-silyl)propionic acid-d4, Aldrich 269913)), centrifuged for 5 minutes at 4°C at 13000 rpm, and then 600 μl was put in a 5 mm short tube. Stools were re-suspended in a cold buffer (0.1 M sodium phosphate pH 7.4, 150 mM NaCl, 10%D2O, 0.05% NaN3) at ratio 1mg:10μL, vortexed for 1 minute, incubated for 15 minutes on ice and centrifuged at 4°C at 13000 rpm, keeping the fecal extract and repeat the protocol extraction twice as reported in Zhiqiang P. et al^111^. 600μL of fecal extracts were mixed with DSS at a final concentration of 0.05 mM and put in a 5 mm short tube. NMR spectra were recorded on a Bruker Avance 600 Ultra Shield TM Plus 600-MHz Spectrometer equipped with a triple-resonance cryoprobe and pulsed field gradients. In addition, the NMR spectrometer is equipped with the SampleJet autosampler. On urine samples,^1^H-1D NMR nuclear Overhauser enhancement spectroscopy (noesypr1d) with a spectral width of 20 ppm, an acquisition time of 2 seconds, a recycle time of 6 seconds, and 32 transients,^1^H-1D NMR Carr-Purcell-Meiboom-Gill (cpmgpr1d) was recorded with an acquisition time of 2 seconds, a recycle time of 4 seconds, and 32 transients were acquired. On fecal extract samples, ^1^H-1D NMR nuclear Overhauser enhancement spectroscopy (noesypr1d) with a spectral width of 20 ppm, an acquisition time of 4 seconds, a recycle time of 6 seconds, and 256 transients were acquired.

### NMR SPECTRA PROCESSING AND MULTIVARIATE ANALYSIS

All acquired spectra were processed with zero filling to 132k points and apodized with an unshifted Gaussian and a 0.5-Hz line broadening exponential using Mnova 14.1.2 (Mestrelab Research S.L.). Spectra were aligned and divided into regular buckets (0.04 ppm), excluding DSS, urea, and water regions, to be subjected to multivariate analysis. Binned spectra were then normalized to total area, scaled (autoscaling for urine spectra and Pareto scaling for feces spectra were applied), and subjected to Partial Least-Squares Discriminant Analysis (PLS-DA) using Metaboanalyst 5.0 [https://www.metaboanalyst.ca/home.xhtml]. Metabolites were selected according to Variable Importance of Projection with a VIP score >1.5. R2 and Q2 were assessed through the cross-validation (CV) method, 5-fold CV.

### NMR METABOLITES IDENTIFICATION AND QUANTIFICATION

Metabolites’ identification and peaks’ assignment were done with the support of literature and comparison with Chenomx NMR Suite version 8.6 (Chenomx, Inc., Edmonton, AB, Canada) and online database Human Metabolome DataBase (HMDB https://hmdb.ca/). The Simple Mixture Analysis (MNova SMA v. 3.0.0.8253) plugin integrated in MNova software was used to set a semi-automatic protocol for the identification and quantification of metabolites. Concentrations were expressed as μmol/mmol for urine samples, normalizing the value for the creatinine, and as mM for fecal extracts. Univariate analysis was performed using Prism (version 10.4.2, Graphpad software). For comparing the HC and the MS groups, the Mann-Whitney test corrected for False Discovery Rate (FDR) was applied. For comparing disease and sex, two-way ANOVA with Tukey correction was applied. Fold change (FC), as the ratio between the average of each metabolite was calculated comparing groups according to disease or healthy state, and disease and sex.

### TARGETED SHORT-CHAIN FATTY ACID PROFILING

Human plasma and CSF samples are analyzed for acetic acid (C2), propionic acid (C3), and butyric acid (C4) by GC-MS. Acetic acid, propionic acid, and butyric acid analytical standards were purchased by Sigma-Aldrich and diluted in water at these final concentrations to make a calibration curve: Acetic acid: 50 μM-100 μM-150 μM-200 μM; Propionic acid: 0.5 μM-1 μM-5 μM-10 μM; Butyric acid: 0.1 uM-0.5 μM-1 μM-2 μM. Metabolites extraction and derivatization: The Protocol for SCFA extraction and derivatization was adapted from previously published protocols^112–114^. 100 μL of SCFA standards, CSF, or plasma were derivatized by adding 40 μL of phosphate buffer 0.1M pH 8, 280 μL of 100 mM pentafluorobenzyl bromide (PFBBr) in acetone, and 1 μL of 500 μM Acetic Acid d3 as internal standard. Samples were vortexed and placed in a thermo-block at 60° C for 1 hour. After cooling down, 200 μL of hexane was added, and samples were vortexed for 3 minutes, followed by centrifugation at 10000 rpm for 5 minutes at 4°C. 100 μL of the upper hexane phase was transferred to a glass insert in a GC-vial for MS analysis.

### GC–MS ANALYSIS

Samples were analyzed by GC-MS using a DB-FATWAX UI (30 m × 0.25 mm x 0.25 μm) installed in an Agilent Intuvo 9000 gas chromatograph (GC) interfaced with a 5977B mass spectrometer (MS) (Agilent Technologies, Santa Clara, CA, USA) operating under electron impact (EI) ionization at 70 eV. The temperatures of the inlet, ion source, and transfer line were 250°C, 290°C, and 230°C, respectively. The GC oven temperature was held at 80 °C for 0.5 min, then ramped to 150 °C at a rate of 10 °C/min, to 240°C at 35°C/min, and then maintained at 240°C for 2 minutes, followed by 1 minute of post-run. Samples (1 μl) were injected in splitless mode using helium as the carrier gas at a flow rate of 1.2 ml/min. Solvent delay was set at 5.5 minutes. MS data were collected in SIM mode and monitored as follows:

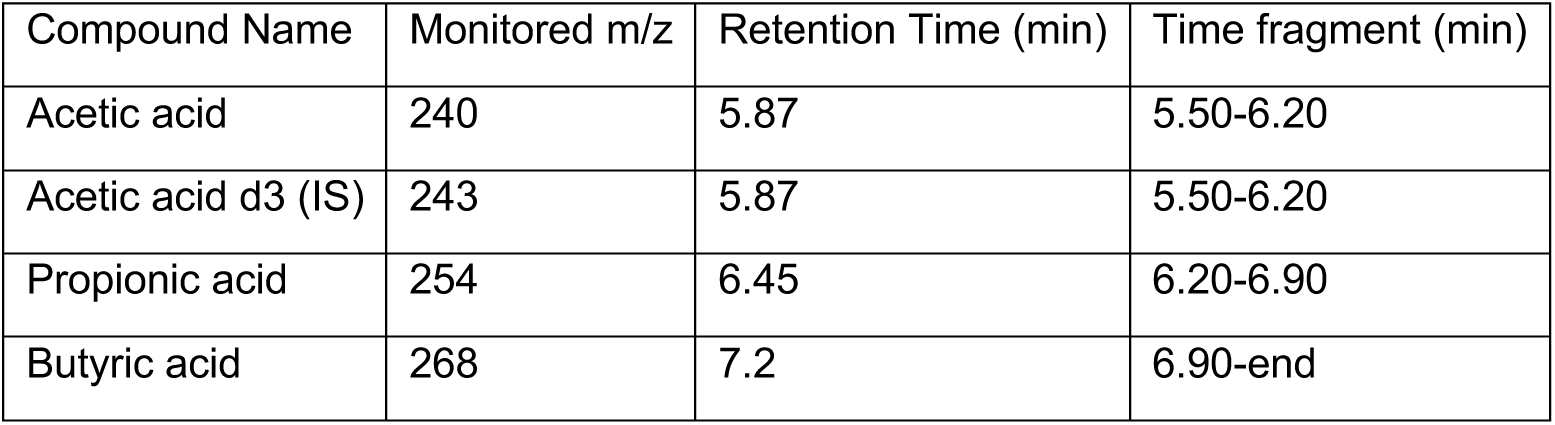

### GC-MS DATA ANALYSIS

Agilent Mass Hunter Quantitative (version 10.2) was used to process the GC–MS data. The concentration of each SCFA in a biological sample was calculated using the calibration curve constructed from the GC–MS data of a corresponding SCFA standard and normalized for the internal standard.

### MEASUREMENTS OF FABP2/i-FABP

Plasma FABP2/i-FABP concentration was determined using the Quantikine anti-Human-iFABP (DFBP20) (R&D Systems Bio-Techne) ELISA kit according to the manufacturer’s instructions. Plasma samples were diluted 5 times in Calibrator Diluent RD5-5 (50μL of plasma sample + 200μL of diluent buffer).

## QUANTIFICATION AND STATISTICAL ANALYSIS

### WHOLE METAGENOME SHOTGUN STATISTICAL ANALYSIS

Statistical analyses and graphical representations were performed in R with packages vegan (v2.6.10)^108^, phyloseq (v1.50.0)^107^, ggplot2 (v3.5.1)^115^, ggpubr (v0.6.0)^116^, gplots (v3.2.0)^117^. Differences in age, sex, BMI, and smoking habits were assessed between the two cohorts with the Fisher test for sex and smoking, the Mann-Whitney test for age, and the unpaired t test for BMI. Differences between the two groups were assessed with Wilcoxon rank-sum tests. Two different types of filters were applied: a) To highlight disease-related microbiota: minimum 20% prevalence, b) To highlight core-microbiota: minimum 80% prevalence and minimum abundance of 0.001. All tests were two-sided except where specified otherwise. Correction for multiple testing (Benjamini–Hochberg procedure, FDR) was applied when appropriate, and significance was defined at Padj < 0.1.

### ADDITIONAL RESOURCES

Clinical Trial registry number: <u>NCT06746896</u>

