## Supplementary material for "Integrated metabolomic and metagenomic profiling reveals distinct signatures in treatment-naïve multiple sclerosis patients": Perego et al_BiorXiV_20251128 supplementary.pdf

### SUPPLEMENTAL INFORMATION

**Document S1. Figures S1–S5 and Tables S1-S3**

**Figure S1. Cohort features (related to Table 1)**

(A) Age and BMI distribution of HC and MS donors, considering the sex component. In the BMI plot, the light grey box indicates the range for normal BMI.

(B) Disease-related features of MS donors: disease duration; disease phase expressed as an arbitrary score (0= remission, 1= relapse or progression); EDSS score. (A-B) Mean + SD, HC F (female HC) n=11; HC M (male HC) n=9; MS F (female MS) n=9; MS M (male MS) n=9. One-way ANOVA, *P < 0.05.

**Figure S2. Prevalence of viral and Blastocystis species in MS compared to HC (related to Fig. 1)**

(A) Pie chart reporting the presence or absence of a specific virus in the microbiota of MS and HC. Numbers represent the raw counts of subjects carrying or not carrying the virus of interest in each category. HC n=20, MS n=17. Only viruses present in at least 20% (8/37) but not in all of the assessed subjects, and with a significant (non-adjusted) p-value (Fisher test), are included in the figure. (B) Table showing viruses present in at least 20% (8/37) but not in all of the assessed subjects, and with a significant (non-adjusted) p-value (Fisher test). Table reporting metadata of subjects positive for Blastocystis; ST1-9: Blastocystis subtypes. 0= absent, 1= present.

**Figure S3. Plasma FABP2 levels do not suggest any detectable gut barrier leakiness in PwMS.**

A-B) Plasma concentrations of FABP2 (pg/ml) in MS and HC were determined by ELISA in donors discriminated only by disease (A), or also by the sex component (B). Comparisons between groups were performed using a two-tailed unpaired t-test for independent samples (A) or one-way ANOVA (B). No statistical significance has been found. HC n=20; MS n= 18. HC F (female HC) n=11; HC M (male HC) n=9; MS F (female MS) n=9; MS M (male MS) n=9.

**Figure S4. District- and sex-specific SCFAs alterations in PwMS (related to Fig. 6).**

(A) FC of mean SCFAs concentration in MS vs HC pooled and sex-specific (male PwMS vs male HC; female PwMS vs female HC). (B) FC of mean CSF SCFAs concentration in female PwMS vs male PwMS. (A-B) Downregulated metabolites (red dots): FC 0-0.7; upregulated metabolites (green dots): FC >1.2. Colored boxes define each metabolite or total SCFAs.

**Figure S5. SCFAs concentration is not correlated with age (related to Fig. 6).**

(A-F) Sex-specific correlations between acetate, butyrate, propionate, and valerate, and the donor's age, were tested by a simple regression model. (A, C, E) Correlation plot with interpolation line in stool (A), plasma (C), and urine (E). (B, D, F) Regression model calculation in stool (B), plasma (D), and urine (F). $r^2$= coefficient of determination; Sy.x= standard error of regression; F= F score; DFn,DFd= degrees of freedom numerator, degrees of freedom denominator. (A-F) HC F (female HC) n=11; HC M (male HC) n=9; MS F (female MS) n=9; MS M (male MS) n=9.

**Table S1. MS donors' clinical features (related to Table 1).**

AST: aspartate transaminase; ALT: alanine transaminase; EDSS: Expanded Disability Status Scale; GERD: gastroesophageal reflux disease; MGUS: monoclonal gammopathy of undetermined significance; LBBB: Left bundle branch block; TBC: tuberculosis; HBV: hepatitis B virus; PPMS: Primary Progressive Multiple Sclerosis; RRMS: Relapsing-remitting multiple sclerosis; BMI: body mass index.

**Table S2. Altered microbiota composition in PwMS compared to HC (related to Fig. 2).**

List of differentially abundant bacteria in PwMS and HC, divided by species, genus, and phylum level. Statistically significant species (Wilcoxon rank-sum test, p-value <0,05, FDR <0,1) are in bold. Species reported in green are more abundant in PwMS, while species written in red are less abundant in PwMS.

**Table S3. Functional analysis of stool microbiota reveals a trend of differentially abundant pathways (related to Fig. 3).**

List of identified functional bacterial microbiome pathways altered in PwMS compared to HC. Pathways with highly significant p-values and lower FDR are in bold (Wilcoxon rank-sum test, p-value <0.05). Pathways reported in green are more abundant in PwMS, while pathways written in red are less abundant in PwMS.

**Table S4. Shotgun metagenomic sequencing**

Read statistics of all samples (number of reads, number of bases, minimum, median, and maximum read length per sample).

**Table S5. NMR metabolomics**

Complete list of metabolites detected by NMR in urine and stool samples.

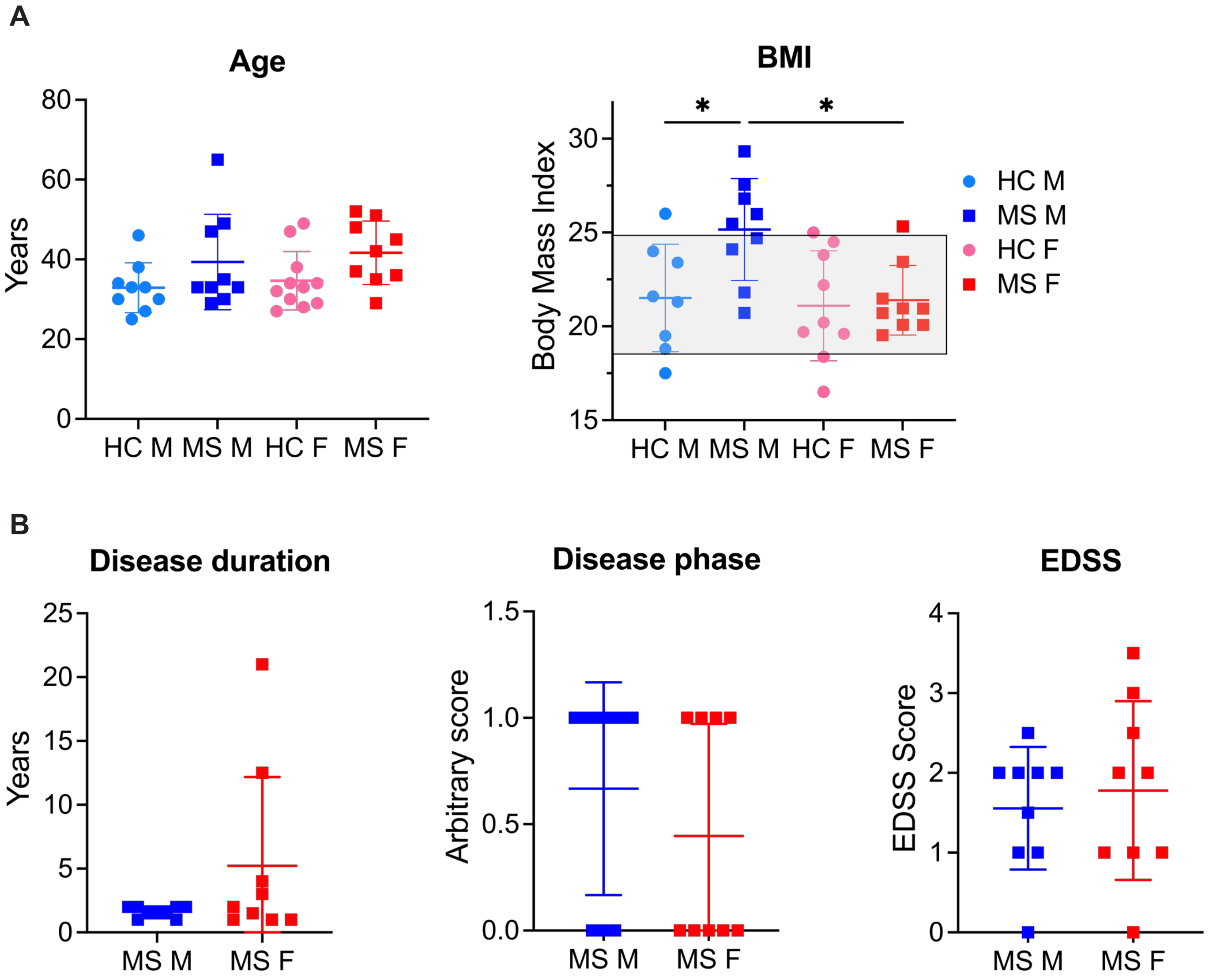
A
Age
Years
80
60
40
20
0
HC M
MS M
HC F
MS F
BMI
Body Mass Index
30
25
20
15
*
*
HC M
MS M
HC F
MS F
B
Disease duration
Years
25
20
15
10
5
0
MS M
MS F
Disease phase
Arbitrary score
1.5
1.0
0.5
0.0
MS M
MS F
EDSS
EDSS Score
4
3
2
1
0
MS M
MS F

**A**

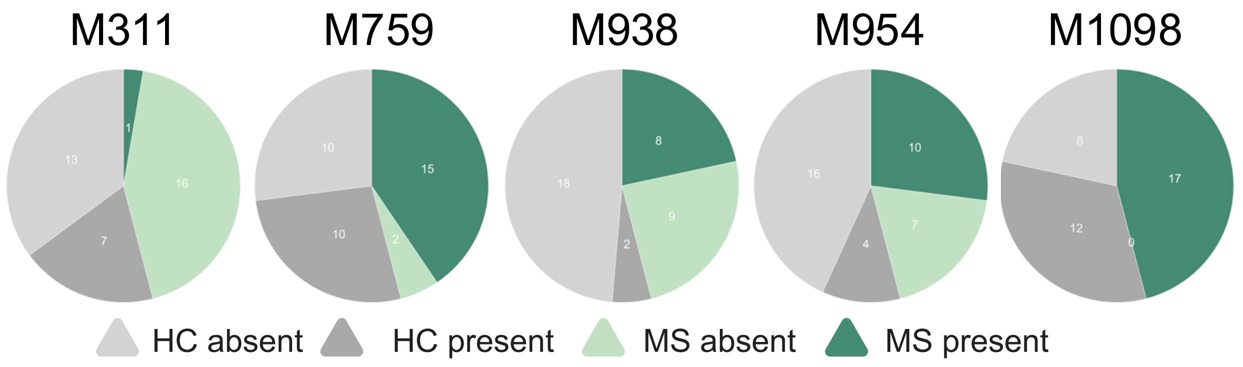


**B**

| *Virus* | *$\chi^2$ p-value* | *Fisher p-value* | *$\chi^2$ p-value adj* | *Fisher p-value adj* | *% HC positive* | *% MS positive* | *Type* | *Genome* |
|---|---|---|---|---|---|---|---|---|
| **M1098** | **0.0035** | **0.0039** | **0.3358** | **0.37** | **60%** | **100%** | **uVSG** | **-** |
| M311 | 0.0510 | 0.0481 | 0.5378 | 0.55 | 35% | 5,8% | kVSG | NC_029014_Parabacteroides_phage_YZ2015b |
| M759 | 0.0170 | 0.0173 | 0.5378 | 0.55 | 50% | 88% | uVSG | - |
| M938 | 0.0250 | 0.0234 | 0.5378 | 0.55 | 10% | 47% | uVSG | - |
| M954 | 0.0175 | 0.0210 | 0.5378 | 0.55 | 20% | 58,8% | uVSG | - |

**C**

| Donor ID | Group | Sex | Age | Diet | ST1 | ST2 | ST3 | ST4 | ST6 | ST7 | ST8 | ST9 |
|---|---|---|---|---|---|---|---|---|---|---|---|---|
| HC4 | HC | M | 33 | Normal diet, no allergies/intollerance | 0 | 0 | 1 | 0 | 0 | 0 | 0 | 0 |
| HC1 | HC | M | 33 | Normal diet, no allergies/intollerance | 0 | 0 | 1 | 0 | 0 | 0 | 0 | 0 |
| HC6 | HC | F | 47 | Normal diet, no allergies/intollerance | 1 | 0 | 0 | 0 | 0 | 0 | 0 | 0 |
| HC5 | HC | F | 49 | Normal diet, no allergies/intollerance | 0 | 0 | 0 | 0 | 1 | 0 | 0 | 1 |

A

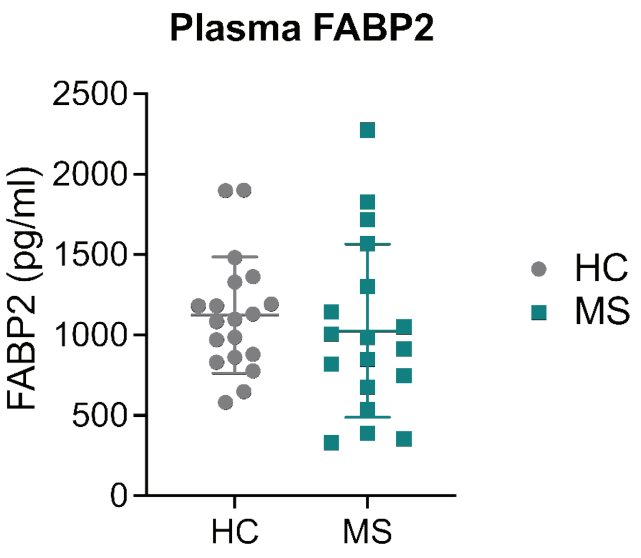
Plasma FABP2
FABP2 (pg/ml)
2500
2000
1500
1000
500
0
HC
MS
HC
MS

B

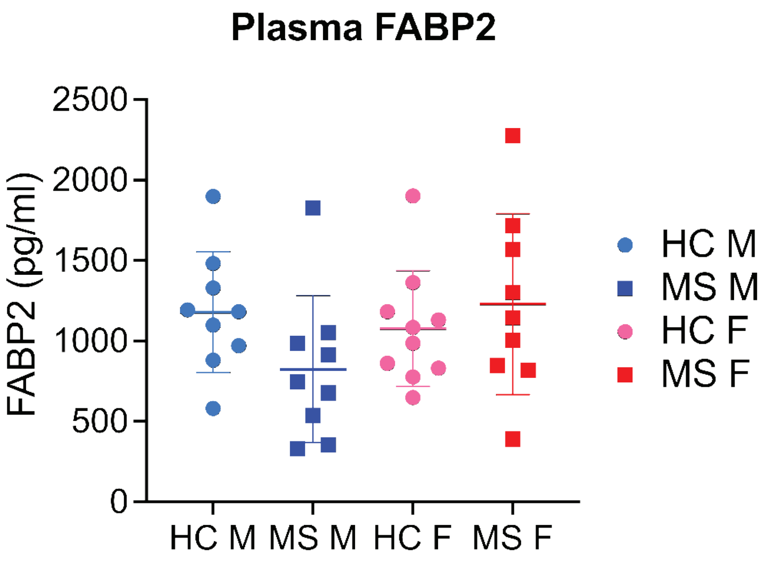
Plasma FABP2
FABP2 (pg/ml)
2500
2000
1500
1000
500
0
HC M
MS M
HC F
MS F
HC M
MS M
HC F
MS F

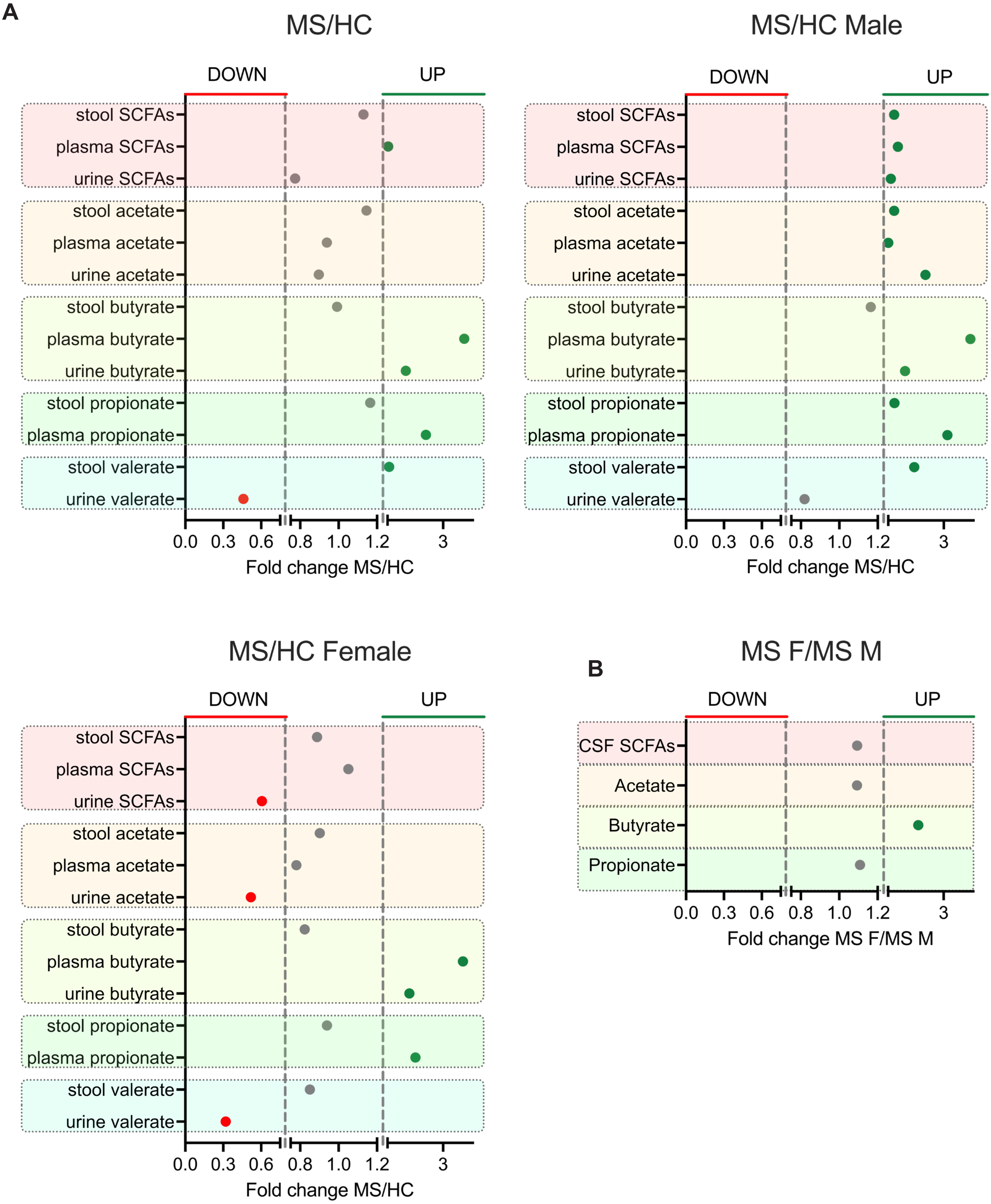
A
MS/HC
DOWN
UP
stool SCFAs
plasma SCFAs
urine SCFAs
stool acetate
plasma acetate
urine acetate
stool butyrate
plasma butyrate
urine butyrate
stool propionate
plasma propionate
stool valerate
urine valerate
0.0 0.3 0.6 0.8 1.0 1.2 3
Fold change MS/HC
MS/HC Male
MS/HC Female
B
MS F/MS M
CSF SCFAs
Acetate
Butyrate
Propionate
Fold change MS F/MS M

**A**

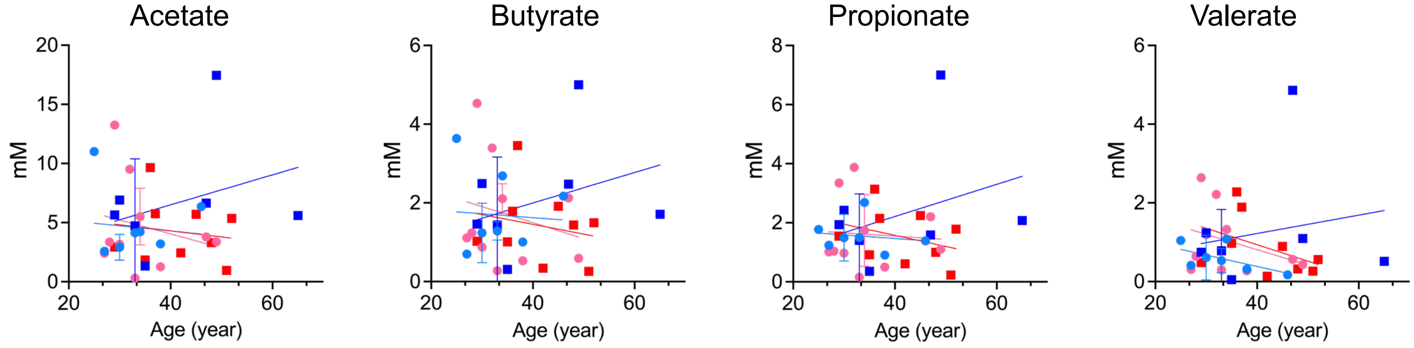


**B**

| | Stool SCFAs | | | | | | | | | | | | | | | |
|---|---|---|---|---|---|---|---|---|---|---|---|---|---|---|---|---|
| | Acetate | | | | Butyrate | | | | Propionate | | | | Valerate | | | |
| | HC M | MS M | HC F | MS F | HC M | MS M | HC F | MS F | HC M | MS M | HC F | MS F | HC M | MS M | HC F | MS F |
| R squared | 0,01025 | 0,08282 | 0,05046 | 0,02322 | 0,003912 | 0,08735 | 0,05093 | 0,04215 | 0,01832 | 0,1002 | 0,004476 | 0,1057 | 0,2255 | 0,03437 | 0,1245 | 0,1948 |
| Sy.x | 2,853 | 5,393 | 3,931 | 2,826 | 1,054 | 1,599 | 1,355 | 1,004 | 0,5948 | 2,086 | 1,275 | 0,9273 | 0,3584 | 1,582 | 0,7982 | 0,7185 |
| F | 0,07246 | 0,6321 | 0,4783 | 0,1664 | 0,02749 | 0,6699 | 0,4829 | 0,308 | 0,1306 | 0,7797 | 0,04047 | 0,827 | 2,038 | 0,2492 | 1,28 | 1,694 |
| DFn, DFd | 1, 7 | 1, 7 | 1, 9 | 1, 7 | 1, 7 | 1, 7 | 1, 9 | 1, 7 | 1, 7 | 1, 7 | 1, 9 | 1, 7 | 1, 7 | 1, 7 | 1, 9 | 1, 7 |
| p-value | 0,7955 | 0,4527 | 0,5066 | 0,6955 | 0,873 | 0,44 | 0,5046 | 0,5962 | 0,7284 | 0,4065 | 0,845 | 0,3934 | 0,1965 | 0,633 | 0,2872 | 0,2343 |
| Deviation from zero? | Non Significant | Non Significant | Non Significant | Non Significant | Non Significant | Non Significant | Non Significant | Non Significant | Non Significant | Non Significant | Non Significant | Non Significant | Non Significant | Non Significant | Non Significant | Non Significant |
| Equation | Y = -0,04342*X + 6,033 | Y = 0,1268*X + 1,432 | Y = -0,1174*X + 8,736 | Y = -0,05134*X + 6,354 | Y = -0,009879*X + 2,019 | Y = 0,03870*X + 0,4495 | Y = -0,04065*X + 3,124 | Y = -0,02482*X + 2,447 | Y = -0,01215*X + 1,952 | Y = 0,05446*X + 0,03284 | Y = -0,01108*X + 1,991 | Y = -0,03756*X + 3,076 | Y = -0,02892*X + 1,541 | Y = 0,02335*X + 0,2875 | Y = -0,03899*X + 2,370 | Y = -0,04165*X + 2,602 |

**C**

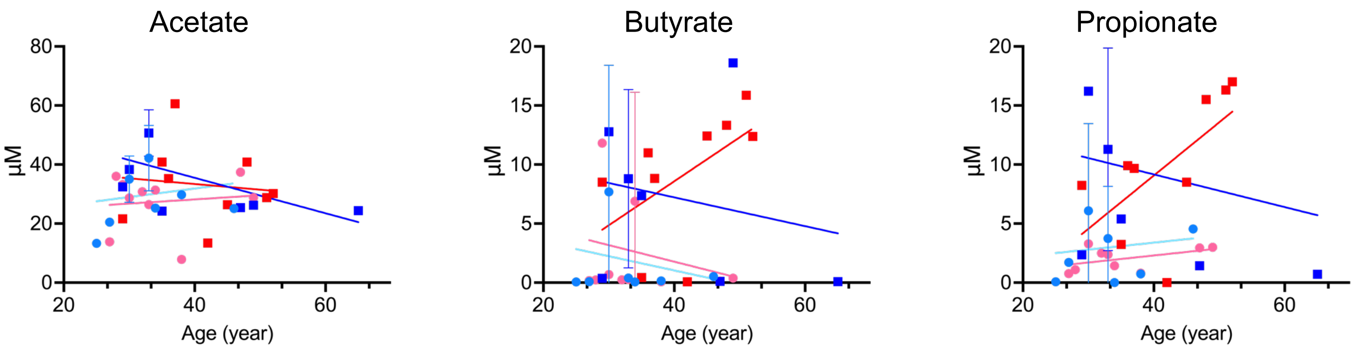


**D**

| | Plasma SCFAs | | | | | | | | | | | |
|---|---|---|---|---|---|---|---|---|---|---|---|---|
| | Acetate | | | | Butyrate | | | | Propionate | | | |
| | HC M | MS M | HC F | MS F | HC M | MS M | HC F | MS F | HC M | MS M | HC F | MS F |
| R squared | 0,02743 | 0,3245 | 0,01331 | 0,01263 | 0,02255 | 0,0395 | 0,04201 | 0,2842 | 0,008983 | 0,03624 | 0,2199 | 0,3793 |
| Sy.x | 11,48 | 11,07 | 9,964 | 14,4 | 5,304 | 7,686 | 5,164 | 5,014 | 4,137 | 9,122 | 0,9449 | 4,894 |
| F | 0,1974 | 3,363 | 0,1079 | 0,08953 | 0,1615 | 0,2879 | 0,3947 | 2,78 | 0,06345 | 0,2632 | 1,974 | 4,277 |
| DFn, DFd | 1, 7 | 1, 7 | 1, 8 | 1, 7 | 1, 7 | 1, 7 | 1, 9 | 1, 7 | 1, 7 | 1, 7 | 1, 7 | 1, 7 |
| p-value | 0,6702 | 0,1093 | 0,751 | 0,7735 | 0,6998 | 0,6082 | 0,5455 | 0,1394 | 0,8084 | 0,6237 | 0,2029 | 0,0774 |
| Deviation from zero? | Not Significant | Not Significant | Not Significant | Not Significant | Not Significant | Not Significant | Not Significant | Not Significant | Not Significant | Not Significant | Not Significant | Not Significant |
| Equation | Y = 0,2884*X + 20,33 | Y= -0,6001*X + 59,51 | Y = 0,1414*X + 22,53 | Y = -0,1919*X + 41,09 | Y = -0,1205*X + 5,853 | Y = -0,1219*X + 12,10 | Y = -0,1401*X + 7,377 | Y = 0,3723*X - 6,312 | Y = 0,05891*X + 1,030 | Y = -0,1384*X + 14,70 | Y = 0,05937*X - 0,07589 | Y = 0,4509*X - 8,965 |

**E**

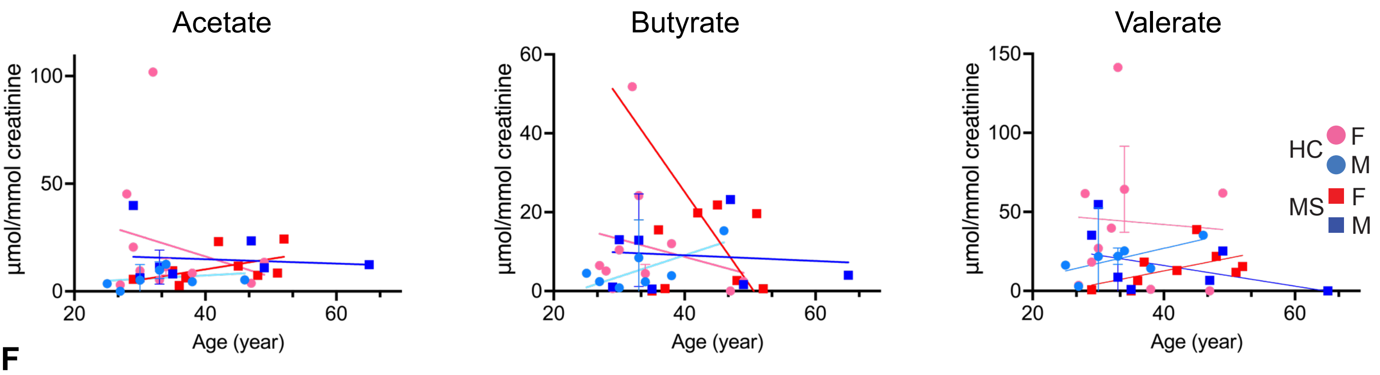


**F**

| | Urine SCFAs | | | | | | | | | | | |
|---|---|---|---|---|---|---|---|---|---|---|---|---|
| | Acetate | | | | Butyrate | | | | Valerate | | | |
| | HC M | MS M | HC F | MS F | HC M | MS M | HC F | MS F | HC M | MS M | HC F | MS F |
| R squared | 0,04462 | 0,0117 | 0,05608 | 0,2576 | 0,007909 | 0,2981 | 0,3382 | 0,04752 | 0,1779 | 0,1598 | 0,003692 | 0,2958 |
| Sy.x | 5,062 | 11,9 | 29,92 | 7,017 | 10,34 | 30,94 | 5,079 | 15,57 | 13,65 | 19,18 | 44,73 | 10,67 |
| F | 0,3269 | 0,08285 | 0,5347 | 2,429 | 0,05581 | 2,974 | 3,578 | 0,4491 | 1,515 | 1,331 | 0,03336 | 2,94 |
| DFn, DFd | 1, 7 | 1, 7 | 1, 9 | 1, 7 | 1, 7 | 1, 7 | 1, 7 | 1, 9 | 1, 7 | 1, 7 | 1, 9 | 1, 7 |
| p-value | 0,5854 | 0,7818 | 0,4832 | 0,1631 | 0,82 | 0,1283 | 0,1005 | 0,5196 | 0,2581 | 0,2865 | 0,8591 | 0,1301 |
| Deviation from zero? | Not Significant | Not Significant | Not Significant | Not Significant | Not Significant | Not Significant | Not Significant | Not Significant | Not Significant | Not Significant | Not Significant | Not Significant |
| Equation | Y = 0,1636*X + 0,8517 | Y = -0,1013*X + 18,98 | Y = -0,9446*X + 53,98 | Y = 0,4871*X - 9,270 | Y = -0,07219*X + 11,97 | Y = -2,376*X + 120,2 | Y = 0,5431*X - 12,64 | Y = -0,4504*X + 26,70 | Y = 0,9496*X - 11,06 | Y = -0,6541*X + 42,26 | Y = -0,3527*X + 56,13 | Y = 0,8149*X - 19,92 |

**Table S1. MS donors' clinical features (related to Table 1).**

| | | | | | | |
|---|---|---|---|---|---|---|
| **disease duration** | 2.0 years | 1 year | 12.5 years | 1.5 years | 4 years | 21 years |
| **comorbidities** | Dysthyroidism, vestibular neuronitis | Hashimoto thyroiditis | NA | Psoriasis | Postpartum depression | NA |
| **clinical phase at diagnosis** | Remission | Relapse | Remission | Remission | Remission | Progression |
| **EDSS at diagnosis** | 1.0 | 2.0 | 3.5 | 2.0 | 1.0 | 3.0 |
| **Oligoclonal bands** | pos | pos | pos | pos | pos | pos |
| **ALT (U/L)** | 12 | 17 | 13 | 17 | 8 | 12 |
| **AST (U/L)** | 15 | 16 | 12 | 21 | 15 | 23 |
| **indirect bilirubin (mg/dL)** | 0.27 | 0.42 | 0.24 | 0.35 | 0.26 | 0.39 |
| **direct bilirubin (mg/dL)** | 0.19 | 0.30 | 0.20 | 0.30 | 0.19 | 0.29 |
| **Total bilirubin (mg/dL)** | 0.46 | 0.72 | 0.44 | 0.65 | 0.45 | 0.68 |
| **Lymphocytes (10*9/L)** | 2 | 2.9 | 2.4 | 2.3 | 1.8 | 2.4 |
| **Neutrophils (10*9/L)** | 2.3 | 3.1 | 4.4 | 2.4 | 1.5 | 2.0 |
| **White blood cells (10*9/L)** | 4.8 | 6.8 | 7.6 | 5.4 | 4.0 | 5.1 |
| **vitamin D (ng/mL)** | 26.7 | 12.0 | 10.6 | 17.1 | 16.1 | 19.8 |
| **Clinical Diagnosis** | RRMS | RRMS | RRMS | RRMS | RRMS | PPMS |
| **BMI** | 23,44 | 20,94 | 20,96 | 20,08 | 20,07 | 20,7 |
| **Age** | 35 | 36 | 45 | 42 | 37 | 51 |
| **Sex** | Female | Female | Female | Female | Female | Female |
| **ID code** | MS1 | MS2 | MS3 | MS4 | MS5 | MS6 |

| | | | | | | | |
|---|---|---|---|---|---|---|---|
| 1 year | 3 years | 1 year | 1.5 years | 1.5 years | 2.0 years | 2 years | 1 year |
| Generalized anxiety disorder, GERD | NA | NA | NA | bronchial asthma, head traumas during sport, ANA | NA | NA | uretrhal stenosis |
| Remission | Relapse | Relapse | Remission | Relapse | Remission | Relapse | Relapse |
| 1.0 | 2.5 | 0.0 | 2.0 | 1.0 | 1.0 | 2.0 | 2.0 |
| pos | pos | pos | pos | neg | pos | pos | neg |
| 18 | 26 | 13 | 9 | 25 | 31 | 12 | 11 |
| 18 | 17 | 18 | 14 | 28 | 24 | 21 | 18 |
| 0.38 | 0.38 | 0.33 | 0.72 | 0.31 | 0.44 | 0.70 | 0.38 |
| 0.28 | 0.21 | 0.22 | 0.43 | 0.20 | 0.25 | 0.32 | 0.21 |
| 0.66 | 0.59 | 0.55 | 1.15 | 0.51 | 0.69 | 1.02 | 0.59 |
| 2.5 | 2.2 | 2.4 | 2.2 | 3 | 2.2 | 2.6 | 1.4 |
| 2,00 | 3.0 | 3.1 | 3.7 | 3.7 | 2.2 | 2.9 | 3.5 |
| 5.4 | 6.0 | 6.1 | 6.5 | 7.6 | 4.52 | 6.5 | 5.4 |
| 26 | S | 15.2 | 16.2 | 12.5 | S (33.7) | 20.2 | 13.2 |
| RRMS | RRMS | RRMS | RRMS | RRMS | RRMS | RRMS | RRMS |
| 21,48 | 25,33 | 19,53 | 20,72 | 26,81 | 24,69 | 21,8 | 25,47 |
| 29 | 52 | 48 | 29 | 47 | 35 | 33 | 49 |
| Female | Female | Female | Male | Male | Male | Male | Male |
| MS7 | MS8 | MS9 | MS10 | MS11 | MS12 | MS13 | MS14 |

| | | | | | | | | | | | | | | | | | | |
|---|---|---|---|---|---|---|---|---|---|---|---|---|---|---|---|---|---|---|
| MS15 | Male | 30 | 25,98 | RRMS | 12.0 | 8.4 | 4.6 | 2.9 | 1.40 | 0.48 | 0.92 | 98 | 248 | pos | 2.0 | Relapse | NA | 2 years |
| MS16 | Male | 33 | 27,55 | RRMS | 14.0 | 8.2 | 3.7 | 3.4 | 0.66 | 0.17 | 0.49 | 28 | 23 | pos | 1.5 | Relapse | Hepatic steatosis | 1 year |
| MS17 | Male | 33 | 29,32 | RRMS | 26.3 | 8.3 | 5,00 | 2.4 | 0.80 | 0.28 | 0.52 | 25 | 36 | pos | 0.0 | Remission | Bipolar disorder | 1.5 years |
| MS18 | Male | 65 | 24,11 | Inflammatory CNS disease | 18.3 | 6.1 | 2.7 | 2.6 | 1.09 | 0.3 | 0.79 | 26 | 24 | neg | 2.5 | Relapse | TBC, MGUS IgG Kappa, Protrusion discal, HBV, Favism, LBBB | 2 years |

AST: aspartate transaminase; ALT: alanine transaminase; EDSS: Expanded Disability Status Scale ; GERD: gastroesophageal reflux disease; MGUS: monoclonal gammopathy of undetermined significance; LBBB:Left bundle branch block; TBC: tuberculosis; HBV: hepatitis B virus ; PPMS: Primary Progressive Multiple Sclerosis; RRMS: Relapsing-remitting multiple sclerosis; BMI: body mass index.

**Table S2. Altered microbiota composition in PwMS compared to HC (related to Fig. 2).**

| *Species* | *p-value* | *FDR* | *r* | *clr median* | *raw median* | *clr median* | *raw median* |
|---|---|---|---|---|---|---|---|
| Roseburia_inulinivorans | 0.0189 | 0.2336 | -0.3858 | 2.3768 | 0.1313 | 0.6195 | 0.0216 |
| Agathobaculum_butyriciproducens | 0.0305 | 0.2336 | -0.3557 | 4.7707 | 0.3558 | 3.6351 | 0.1107 |
| Ruthenibacterium_lactatiformans | 0.0329 | 0.2336 | 0.3507 | 1.1723 | 0.0348 | 2.3027 | 0.0987 |
| Alistipes_communis | 0.0329 | 0.2336 | -0.3507 | 2.5583 | 0.1176 | 1.5157 | 0.0433 |
| Blautia_glucerasea | 0.0355 | 0.2336 | -0.3457 | -3.9254 | 0.1892 | -9.2786 | 0.0666 |
| Faecalibacterium_prausnitzii | 0.0355 | 0.2336 | -0.3457 | 27.6743 | 6.7385 | 20.4744 | 3.8003 |
| Eggerthella_lenta | 0.0355 | 0.2336 | 0.3457 | 1.1232 | 0.0347 | 2.3537 | 0.1172 |
| Ruminococcus_torques | 0.0382 | 0.2336 | 0.3407 | 1.8656 | 0.0761 | 3.4899 | 0.2938 |
| Roseburia_hominis | 0.0382 | 0.2336 | -0.3407 | -0.0392 | 0.0409 | -1.5968 | 0.0105 |
| Flavonifractor_plautii | 0.0443 | 0.2343 | 0.3307 | 1.3762 | 0.038 | 2.6244 | 0.1603 |
| Clostridiales_bacterium_KLE1615 | 0.0476 | 0.2343 | -0.3257 | 3.2115 | 0.2609 | 0.6122 | 0.0175 |
| *Genus level* | | | | | | | |
| **Gordonibacter** | **0.0023** | **0.064*** | **0.501** | **-1.816** | **0.0143** | **1.4441** | **0.0834** |
| **Roseburia** | **0.0031** | **0.064*** | **-0.486** | **1.1096** | **1.6982** | **-4.8917** | **0.2617** |
| **Flavonifractor** | **0.0042** | **0.064*** | **0.471** | **0.3981** | **0.038** | **2.3916** | **0.1946** |
| **Lachnospira** | **0.0088** | **0.1008** | **-0.4309** | **5.4058** | **0.3403** | **-0.4275** | **0.0253** |
| Clostridiaceae_unclassified | 0.0174 | 0.1352 | -0.3908 | -3.4176 | 0.5964 | -7.7207 | 0.2049 |
| Fusicatenibacter | 0.0205 | 0.1352 | -0.3808 | 6.654 | 1.3597 | 5.5327 | 0.504 |
| Eubacterium | 0.0261 | 0.1352 | -0.3657 | -0.5852 | 0.3242 | -5.7634 | 0.1774 |
| Lachnospiraceae_unclassified | 0.0282 | 0.1352 | -0.3607 | -1.722 | 1.5355 | -6.2633 | 0.851 |
| Eggerthella | 0.0305 | 0.1352 | 0.3557 | 0.854 | 0.0349 | 2.0546 | 0.1172 |
| Ruthenibacterium | 0.0329 | 0.1352 | 0.3507 | 1.1723 | 0.0348 | 2.3027 | 0.0987 |
| Faecalibacterium | 0.0329 | 0.1352 | -0.3507 | 29.1383 | 7.8985 | 20.5739 | 4.185 |
| Coprococcus | 0.0382 | 0.1352 | -0.3407 | 17.5248 | 1.5898 | 14.8452 | 0.703 |
| Mediterraneibacter | 0.0412 | 0.1352 | 0.3357 | -8.1449 | 0.8047 | -0.963 | 1.841 |
| Alistipes | 0.0412 | 0.1352 | 0.3357 | 16.9655 | 1.3974 | 23.1265 | 1.8345 |
| Agathobaculum | 0.0443 | 0.1358 | -0.3307 | 6.1236 | 0.3558 | 4.9326 | 0.1107 |
| *Phylum level* | | | | | | | |
| **Firmicutes** | **0.0205** | **0.0658*** | **-0.3808** | **-120.6564** | **61.7137** | **-165.4409** | **54.7176** |
| **Actinobacteria** | **0.0329** | **0.0658*** | **0.3507** | **24.085** | **7.8854** | **43.1878** | **15.2486** |

List of differentially abundant bacteria in PwMS and HC, divided by species, genus, and phylum level. Statistically significant species (Wilcoxon rank-sum test, p-value <0,05, FDR <0,1) are in bold. Species reported in green are more abundant in PwMS, while species written in red are less abundant in PwMS.

**Table S3. Functional analysis of stool microbiota reveals a trend of differentially abundant pathways (related to Fig. 3).**

| | | | | HC | | MS | |
|---|---|---|---|---|---|---|---|
| *Pathway* | *r* | *p-value* | *FDR* | *Clr median* | *Raw median* | *Clr median* | *Raw median* |
| **PWY-5972 stearate biosynthesis I (animals)** | **0.54** | **0.00** | **0.38** | **-3.60** | **0.00** | **-3.14** | **23.80** |
| **PWY-5464 superpathway of cytosolic glycolysis (plants), pyruvate dehydrogenase and TCA cycle** | **0.51** | **0.00** | **0.38** | **-4.21** | **0.00** | **-3.20** | **24.12** |
| **PWY-5677 succinate fermentation to butanoate** | **0.43** | **0.01** | **0.72** | **-3.28** | **18.74** | **-1.77** | **147.67** |
| GLYCOLYSIS-TCA-GLYOX-BYPASS superpathway of glycolysis, pyruvate dehydrogenase, TCA, and glyoxylate bypass | 0.42 | 0.01 | 0.72 | -4.37 | 0.00 | -3.29 | 22.06 |
| PWY66-391 fatty acid β-oxidation VI (mammalian peroxisome) | 0.41 | 0.01 | 0.72 | -2.37 | 66.17 | -1.00 | 299.63 |
| PWY-6895 superpathway of thiamine diphosphate biosynthesis II | -0.41 | 0.01 | 0.72 | 1.51 | 2925.32 | 1.10 | 2114.25 |
| KDO-NAGLIPASYN-PWY superpathway of (Kdo)2-lipid A biosynthesis | 0.40 | 0.01 | 0.72 | -3.48 | 0.00 | -3.13 | 30.56 |
| METHGLYUT-PWY superpathway of methylglyoxal degradation | 0.40 | 0.01 | 0.72 | -2.00 | 55.35 | -1.38 | 165.25 |

| | | | | | | | |
|---|---|---|---|---|---|---|---|
| PWY-5705 allantoin degradation to glyoxylate III | 0.39 | 0.02 | 0.72 | -4.99 | 0.00 | -3.31 | 22.11 |
| FERMENTATION-PWY mixed acid fermentation | -0.38 | 0.02 | 0.72 | 1.86 | 5122.82 | 1.48 | 3910.48 |
| PWY-7094 fatty acid salvage | 0.37 | 0.02 | 0.72 | -4.79 | 3.28 | -2.74 | 45.17 |
| PWY-6876 isopropanol biosynthesis (engineered) | 0.37 | 0.03 | 0.72 | -3.24 | 25.34 | -2.38 | 49.88 |
| GLUCOSE1PMETAB-PWY glucose and glucose-1-phosphate degradation | -0.36 | 0.03 | 0.72 | 2.91 | 9861.37 | 2.32 | 8106.90 |
| PWY-621 sucrose degradation III (sucrose invertase) | -0.35 | 0.03 | 0.72 | 1.81 | 3569.34 | 1.19 | 2914.79 |
| DENOVOPURINE2-PWY superpathway of purine nucleotides de novo biosynthesis II | 0.35 | 0.04 | 0.72 | -1.72 | 0.00 | -1.06 | 167.77 |
| PRPP-PWY superpathway of histidine, purine, and pyrimidine biosynthesis | 0.35 | 0.04 | 0.72 | -1.33 | 0.00 | -0.60 | 267.17 |

| | | | | | | | |
|---|---|---|---|---|---|---|---|
| PWY-7345 superpathway of anaerobic sucrose degradation | -0.34 | 0.04 | 0.72 | 1.78 | 4162.26 | 1.52 | 3693.80 |
| P221-PWY octane oxidation | 0.34 | 0.04 | 0.72 | -5.42 | 2.98 | -3.49 | 28.47 |
| PWY-6992 1,5-anhydrofructose degradation | 0.34 | 0.04 | 0.72 | -4.81 | 0.00 | -2.85 | 18.16 |
| PWY0-1338 polymyxin resistance | 0.33 | 0.04 | 0.72 | -4.32 | 0.00 | -2.77 | 58.34 |
| PWY0-1337 oleate β-oxidation | 0.33 | 0.05 | 0.72 | -4.58 | 2.28 | -2.68 | 45.87 |

List of identified functional bacterial microbiome pathway altered in PwMS compared to HC. Pathways with highly significant p-value and lower FDR are in bold (Wilcoxon rank-sum test, p-value <0.05). Pathways reported in green are more abundant in PwMS, while pathways written in red less abundant in PwMS.
